# Species better track the shifting isotherms in the oceans than on lands

**DOI:** 10.1101/765776

**Authors:** J. Lenoir, R. Bertrand, L. Comte, L. Bourgeaud, T. Hattab, J. Murienne, G. Grenouillet

## Abstract

Despite mounting evidence of species redistribution as climate warms, our knowledge of the coupling between species range shifts and isotherm shifts is limited. Compiling a global geo-database of 30,534 range shifts from 12,415 taxa, we show that only marine taxa closely track the shifting isotherms. In the oceans, the velocity of isotherm shifts interacts synergistically with anthropogenic disturbances and baseline temperatures such that isotherm tracking by marine life happens either in warm and undisturbed waters (e.g. Central Pacific Basin) or in colder waters where human activities are more pronounced (e.g. North Sea). On lands, increasing anthropogenic activities and temperatures negatively impact the capacity of terrestrial taxa to track isotherm shifts in latitude and elevation, respectively. This suggests that terrestrial taxa are lagging behind the shifting isotherms, most likely due to their wider thermal safety margin, more constrained physical environment for dispersal and higher availability of thermal microrefugia at shorter distances.

---

The redistribution of life on Earth in response to climate change (*1–4*) is now considered as a global change driver on its own with important consequences on ecosystems and human well-being (*5*). As climate warms, isotherms are shifting poleward to cooler latitudes and upslope to colder elevations, generating spatially-structured patterns in the velocity of isotherm shifts (*6, 7*). Marine taxa seem to closely track this complex mosaic of climate velocities (*8*). Yet, the pattern is less clear for terrestrial organisms (*2*). Evidence suggests that biotic responses on lands are lagging behind the velocity of climate change, particularly for long-lived taxa and poor-dispersers (*9, 10*). To date, a comprehensive analysis of the coupling between the velocity of species range shifts and the velocity of isotherm shifts across biological systems and life forms is still lacking.

To fill this knowledge gap, we compiled range shifts for marine and terrestrial taxa from an exhaustive literature review building on and updating the most recent syntheses on climate-related range shifts (see section Materials and Methods below). Based on this review, we compiled a geo-database (Table S1 and Figs. S1-S2 in the Supplementary Materials) of 30,534 range shifts covering four kingdoms (*Bacteria, Plantae, Fungi* and *Animalia*), 20 phyla, 56 taxonomic classes and 12,415 taxa (Fig. S3). Focusing on latitudinal and elevational gradients, we first provided robust estimates of the velocity of range shift for the 20 most studied taxonomic classes (Fig. 1) by fitting models accounting for different methodological attributes among studies (*11*) (Fig. S4). Models were arranged in a full factorial design of geographical gradient (latitude *vs*. elevation) × biological system (marine *vs*. terrestrial) × hemisphere (North *vs*. South) × positional parameter (centroid *vs*. margins) (Table S2). Then, we assessed the coupling between the velocity of isotherm shifts and the velocity of range shifts at the species level, separately for the marine and terrestrial realm and/or the latitudinal and elevational gradient. As before, we controlled for varying methodologies and tested for two-way interaction terms between the velocity of isotherm shifts and life-form categories (ectotherms, endotherms, phanerogams, cryptogams), mean annual temperature conditions prior to the baseline surveys (baseline temperatures) and the human footprint index (*11, 12, 13*).

**Fig. 1.**
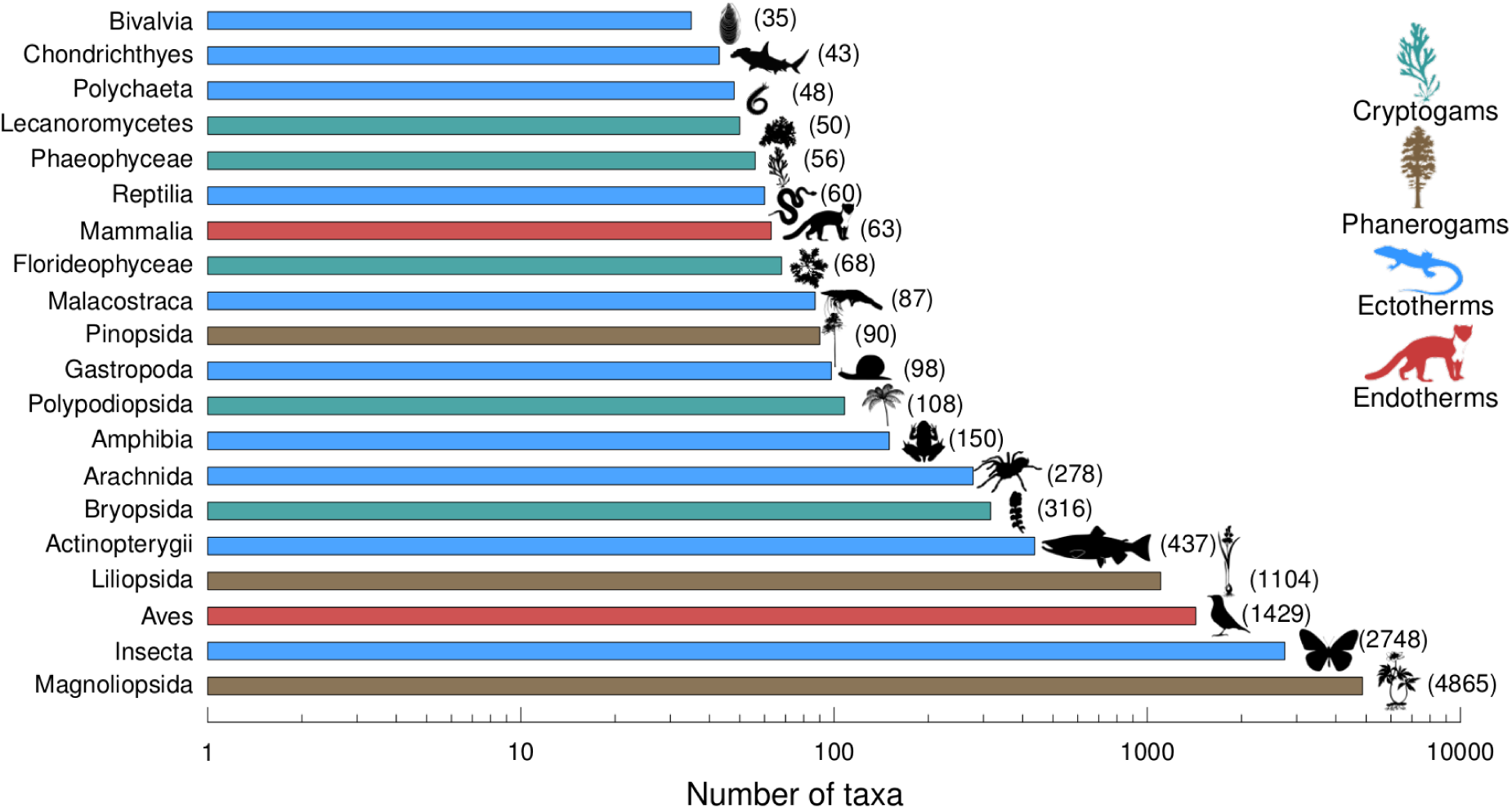
Number of taxa, in log scale, per taxonomic class: from the least (top) to the most (bottom) studied taxonomic class. Only taxonomic classes with more than 30 observations per factorial model are displayed (N = 20; Table S3) (see Fig. S3 for full taxonomic coverage). The exact number of taxa per taxonomic class is provided in parenthesis.

We found a large variation in the estimated velocity of range shift per taxonomic class, ranging from 0.70 m.yr^−1^ upslope for fish to 18.54 km.yr^−1^ upward for insects (Fig. 2; Figs. S5-S7; Table S3). Our models explained from 6 to 97% (mean = 46%) of the total variation in the velocity of range shifts, of which methodological attributes contributed from 6 to 82% (mean = 36%) (Table S2). However, we found no evidence suggesting that single-species studies are more likely to show larger poleward shifts in latitude (Fig. S4B) or larger upward shifts in elevation (Fig. S4C) than studies reporting range shifts for multiple species (*1*). This confirms the importance of accounting for varying methodologies in quantitative reviews (*11*) and points to the underutilized potential of single-taxon studies to our understanding of biological responses to climate change.

**Fig. 2.**
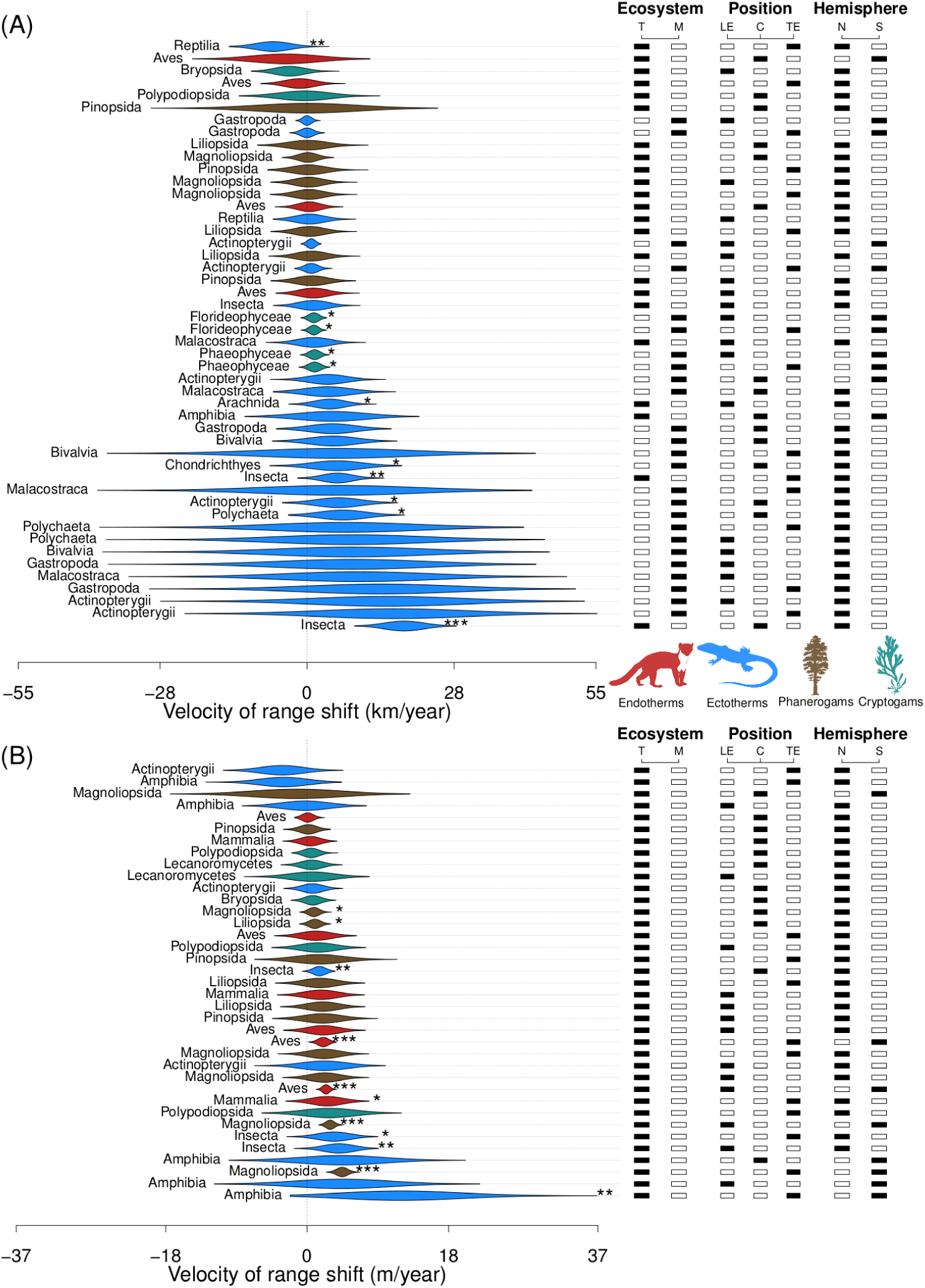
Estimated velocity of range shift per taxonomic class (i.e. effect size) in km.yr^−1^ and m.yr^−1^ for latitudinal (**A**) and elevational (**B**) range shifts, respectively. Outputs are displayed for all possible combinations of positional parameter (TE: trailing edge *vs*. CE: centroid *vs*. LE: leading edge) × hemisphere (N: north *vs*. S: south) × biological systems (M: marine *v*s. T: terrestrial) (see Figs. S5-S7 for a different display by factorial model). Violin plots represent the distribution of 5,000 bootstrap iterations. Stars show significant deviations from zero shift (*: *P* < 0.05; **: *P* < 0.01; ***: *P* < 0.001).

Marine taxa (∼80% being ectotherms in the database; Fig. S2C) have moved towards the poles at a mean (±s.e.m.) pace of 5.92±0.94 km.yr^−1^, almost six times faster than terrestrial taxa (1.11±0.96) (one-way ANOVA: F = 12.68; *df* factor = 1; *df* residuals = 45; *P* = 0.0009). These rates far exceed the mean latitudinal range shift (0.61±0.24) reported by the seminal synthesis (*1*). Yet, we found lower velocities than the values reported by subsequent syntheses focusing chiefly on terrestrial taxa (1.76±0.29) (*2*) or exclusively on marine taxa (7.20±1.35) (*3*). Contrarily to a previous report (*3*) but according to expectations from thermal tolerance limits of marine ectotherms (*14*), our results suggest that both the leading (6.02±1.77) and trailing (6.49±2.13) edge of marine taxa are equally responsive to warming (one-way ANOVA: F = 0.15; *df* factor = 2; *df* residuals = 21; *P* = 0.86). Similarly, along the elevational gradient, we found that terrestrial taxa have moved towards the summits at a mean pace of 1.78±0.41 m.yr^−1^ (*P* = 2.7×10^−5^), irrespective of the range position (one-way ANOVA: F = 1.38; *df* factor = 2; *df* residuals = 34; *P* = 0.27). This supports a recent synthesis concluding on symmetric boundary shifts in mountains (*15*). However, it does not necessarily imply that all species, irrespective of their surrounding environments and intrinsic characteristics, are equally responsive to isotherm shifts (Figs. S8-S10).

Unlike terrestrial ectotherms, marine ectotherms closely track isotherm shifts in latitude, the extent to which being conditional on experienced temperature regimes and anthropogenic pressures (variance explained = 33%; Tables S4-S5; Fig. 3) (*8*). We found similar results for marine cryptogams (Fig. S11). More specifically, the velocity of isotherm shifts interacts synergistically with the standardized human footprint index (e.g. fishing pressure) and baseline temperatures (Table S5) (Figs. 3C-D), such that isotherm tracking by marine ectotherms and cryptogams happens either in initially warm and undisturbed waters (e.g. Central Pacific Basin) (*12*) or in initially colder waters where human activities are more pronounced (e.g. North Sea) (Fig. 4 and Fig. S12). This may stem from the combination of two processes. First, marine ectotherms are living closer to their upper thermal limits in the tropics, where sea surface temperatures are the highest, thus increasing the likelihood of local extirpations at their trailing edges as climate warms (*16*). Second, greater dispersal and colonization abilities in the oceans (*3*) may help marine ectotherms and cryptogams to expand their distribution towards the newly available habitats at their leading edge. By contrast, at high latitudes, where the thermal safety margin of marine ectotherms is larger (*16*), climate warming alone is unlikely to explain isotherm tracking. Instead, anthropogenic activities (e.g. fishing pressure in the North Sea) may render populations more sensitive to climate change at the trailing edge of the range by reducing abundance and density, truncating the age distribution and leading to the depletion of fish stock from previous stronghold (*17*). In parallel, successful management actions at high latitudes in the northern hemisphere, combined with climate warming, may increase the population size of commercial fish at the leading edge of their range (*18*).

**Fig. 3.**
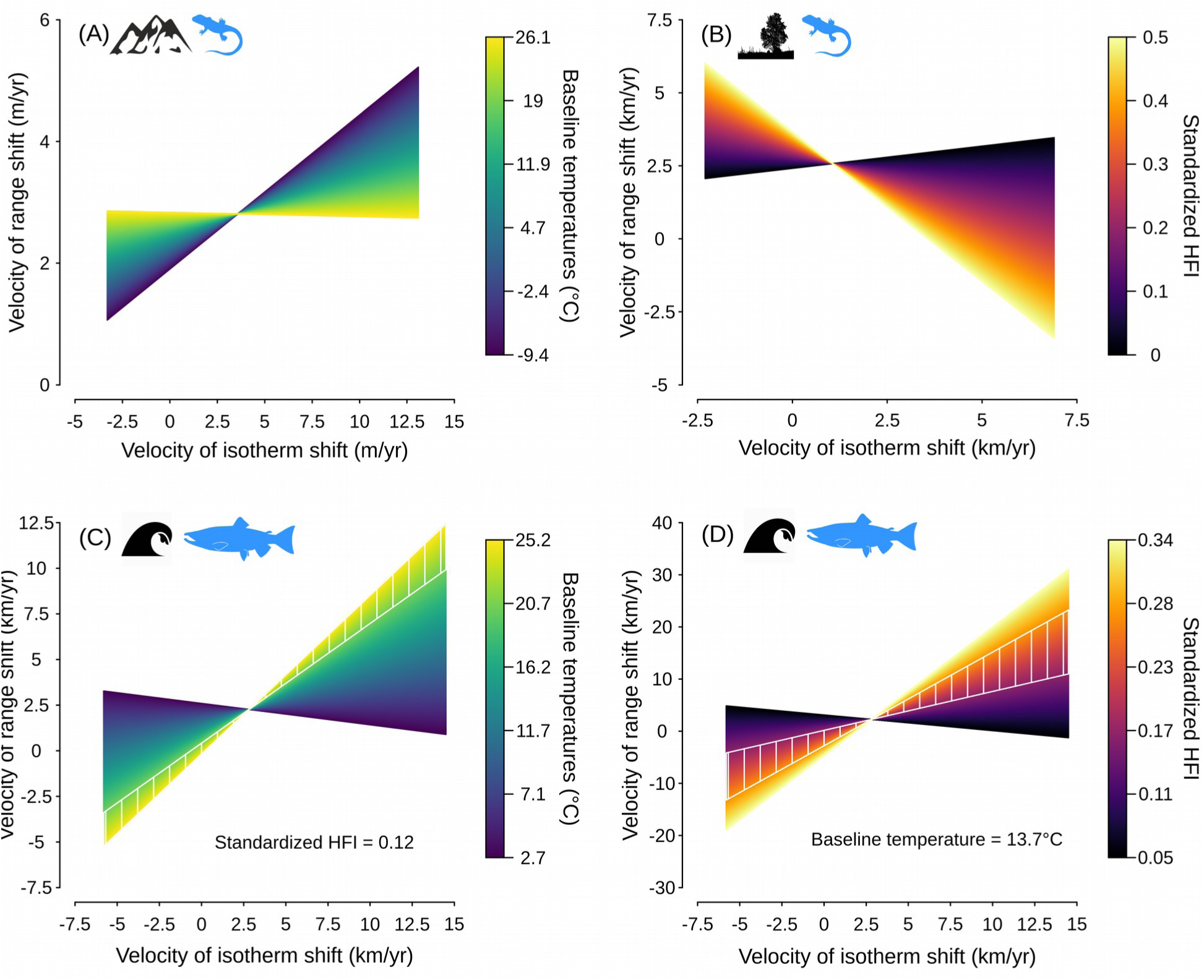
Main determinants of the velocity of species range shifts along the (**A**) elevational and (**B, C, D**) latitudinal gradients for both the (**A, B**) terrestrial and (**C, D**) marine realm. Panel **A** shows the interaction effect between baseline temperatures and the velocity of isotherm shifts in elevation for ectotherms (see. Fig. S14 for the other life forms). Panel **B** shows the interaction effect between the standardized human footprint index and the velocity of isotherm shifts in latitude for terrestrial ectotherms. Note that the same pattern is observed for the three other studied life forms of the terrestrial realm, but the intercept is lower (see Table S5 and Fig. S13). Panel **C** shows the interaction effect between baseline temperatures and the velocity of isotherm shifts in latitude for marine ectotherms while setting the standardized human footprint index to its median value in the database (see. Fig. S11 for marine cryptogams). Panel **D** shows the interaction effect between the standardized human footprint index and the velocity of isotherm shifts in latitude for marine ectotherms while setting baseline temperatures to the median value in the database (see. Fig. S11 for marine cryptogams). The two white lines and the white hatching represent the range of conditions for which marine ectotherms closely track the shifting isotherms in latitude (i.e. slope parameter not significantly different from 1 based on 5,000 bootstrap iterations).

**Fig. 4.**
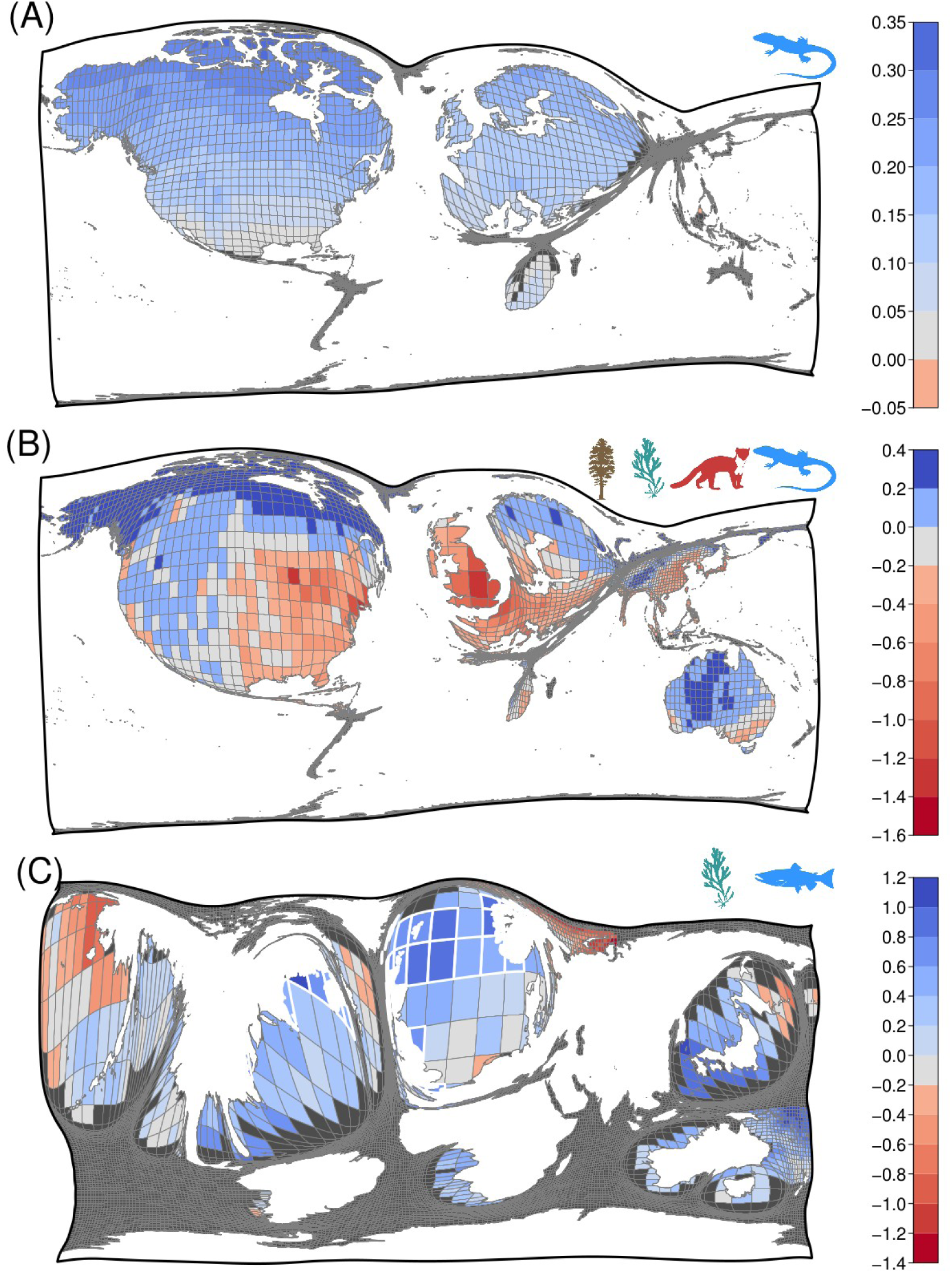
Cartograms of the predicted slope coefficient between the velocity of species range shifts and the velocity of isotherm shifts per 2° × 2° grid cell along the (**A**) elevational and (**B, C**) latitudinal gradient for both the terrestrial (**B**) and (**C**) marine realm. Note than panel **A** only displays the predicted slope coefficient for ectotherms (see Fig. S15 for the three other life forms). Positive slope values (bluish colors) close to 1 suggest a perfect isotherm tracking while negative values (reddish colors) suggest that species are not tracking the shifting isotherms. The number of range shift estimates (i.e. sample size) in each grid cell was used to distort the map: the bigger the grid cell, the larger the sample size (see Fig. S1). Grid cells with a white and bold border display areas where species are closely tracking the shifting isotherms (i.e. slope parameter not significantly different from 1 based on 5,000 bootstrap iterations).

On lands, we found a negative impact of human activities on the capacity of species to track isotherm shifts in latitude (variance explained = 47%; Tables S4-S5; Fig. 3B; Fig. S13). This climatic debt (*10, 19*) suggests that habitat loss and fragmentation (*20*), combined with poor dispersal abilities (*21*), impede the capacity of terrestrial taxa to track isotherm shifts at the leading edge of their range. This, in turn, confirms that isotherm tracking is very unlikely for terrestrial taxa living in the lowlands (*9, 19*). High levels of habitat loss and fragmentation may even outweigh the positive impact that climate warming may have on population dynamics at the leading edge of a species range (*20*) and could explain why some species are shifting in the opposite direction to what is expected based solely on isotherm shifts (Fig. 3B and Fig. S13). In addition, air conducts heat 25 times less effectively than water, which makes terrestrial taxa, in general, less sensitive than marine taxa to temperature fluctuations and thus less likely to move as a direct response to climate warming (*14*). Terrestrial ectotherms may also access thermal microrefugia (e.g. shaded environments) more easily to regulate their body temperature, increasing their thermal safety margin by 3°C, on average, as compared with marine ectotherms (*16*). Therefore, terrestrial ectotherms could better withstand the increase in temperatures and avoid extirpation at the trailing edge of their latitudinal range by staying put or by finding thermal microrefugia at relatively short distances. This is in agreement with our results for mountainous ecosystems, where terrestrial ectotherms seem to better track isotherms upslope, especially under initially cold conditions, than endotherms, phanerogams or cryptogams (variance explained = 11%; Tables S4-S5; Figs. 3A-4A; Figs. S14-S15).

To conclude, the coupling between species range shifts and isotherm shifts is not uniform across realms, confirming the lags observed in the biotic responses of terrestrial organisms to climate change (*9, 10*). Noteworthy, we demonstrate complex interactions between the velocity of climate warming, the human footprint index, temperatures and species characteristics. These interactions need to be integrated into future models to improve scenarios of biodiversity redistribution and its consequences on human well-being (*5*) under future climate change. This said, it is important to bear in mind that our findings, as well as former syntheses on the topic, are still dependent on data availability and thus suffer from severe taxonomic (Fig. 1; Fig. S3) and geographic (Fig. 3; Figs. S1-S2) biases. These limitations may affect our perception of species redistribution, and by consequence challenge global biodiversity conservation efforts (*4, 22*).

## Materials and Methods

### Literature search

We reviewed the scientific and peer-reviewed literature reporting climate-driven range shifts under contemporary climate change. By contemporary climate change, we here mean the period stretching from the beginning of the 19th century and onwards. As a general approach, we started from the reference lists of the most recent meta-analyses and syntheses on the topic (*2–4*) that we completed by regularly searching the scientific literature published between 2014 and 2018 following the same protocol as in Lenoir & Svenning (*4*) (see Appendix 1 therein for the detailed approach: list of keywords and search terms used as well as the list of search engines queried). Because of the clear focus on latitudinal and elevational range shifts in the scientific literature and the lack of information on the other geographical dimensions (*4*), we excluded several reports focusing exclusively on bathymetric or longitudinal range shifts. Broad inclusion criteria comprised studies: (i) focusing on relatively recent (since 1850s) distribution changes; (ii) based on occurrence or abundance data of at least one non-introduced taxon; and (iii) only if studies were based on assessments covering at least two historical censuses. Hence, we excluded studies reporting distribution changes from a single census (synchronous approach) or based on historical patterns of species mortality obtained from climatic reconstructions only, without real occurrence or abundance data to confirm model outputs. Data were only included in our database if the range shift estimates were reported at the taxa (i.e. subspecies, species or genus) level in the original research. When data on species range shifts were not directly available in the text or in tables in the main text or in the appendices of the publication, we first contacted the corresponding authors and requested the data. In case of no positive response, we used the “WebPlotDigitizer” program (https://automeris.io/WebPlotDigitizer/) to extract range shift values from figures, when possible. This drastic selection procedure to ensure data quality led to a total of 30,534 range shift estimates at the taxa level extracted from 258 publications (Table S1). When range shifts were reported for more than one geographically distinct survey area or more than one period delimited by two censuses (e.g. more than one resurvey of historical data), we considered them as independent case studies (N = 325). For each of these 325 case studies, we digitized the study region in Google Earth and used the resulting polygons to retrieve spatial information such as the total area covered by the study. If no clear maps delineating the study area was reported in the original study (e.g. map displaying the study region), we used national geographic boundaries or any meaningful spatial information from the text to delineate the study area. All spatial polygons were used to produce a geo-database (Figs. S1-S2).

Range shift estimates, as reported by the original authors, were coded as positive values if poleward in latitude, or upward in elevation, and negative otherwise (equatorward and downward). When the authors reported horizontal range shifts with both the magnitude and direction (i.e. azimuth) values, we used trigonometric relationships to transform these values into latitudinal range shifts for consistency with the main bulk of data available in the scientific literature. Next, we divided each range shift value by the study duration between two consecutive censuses (ending year – starting year + 1) to assess the rate of range shift (*ShiftR*), or the velocity of biological shifts, in kilometer per year along the latitudinal gradient and in meter per year along the elevational gradient. In addition to the rate or velocity of range shift at the taxa level, we also retrieved information at the case study level, including: the size of the study area (*Area*); the starting year of the study (*Start*); the ending year of the study (*End*); as well as several other methodological attributes known to potentially affect the velocity of range shift (*11*). More specifically, we considered: (i) the number of taxa in a study (*Ntaxa*) (continuous variable ranging from 1 to 4426; median = 21; mean = 122); (ii) the frequency of sampling (*Sampling*) (factor variable with four levels: “continuous”; “irregular”; or comparison of “two”; or “multiple” periods); (iii) whether range shift estimates were generated from “occurrence” or “abundance” data (*PrAb*) (factor variable with two levels); (iv) the spatial resolution of the raw data used to estimate range shifts (*Grain*) (factor variable with three levels: “fine” for data based on GPS coordinates with a spatial resolution lower than 10 km; “coarse” for data based on range maps or atlas grids with a spatial resolution greater than 100 km; and “medium” for intermediate situations); (v) the quality of the approach used to estimate range shifts (*Quality*) (factor variable with four levels: “low” when no data cleaning procedures were performed before computing range shifts; “balanced” when data cleaning or resampling procedure were carried out to calculate range shifts on a balanced dataset; “modeled” when range shifts were obtained from model outputs; and “resurveyed” when range shifts were calculated from paired designs such as permanent plots); and (vi) whether the “significance” of range shift estimates were assessed or “not” in the original study (*Signif*) (factor variable with two levels). To improve the balance in the number of observations among levels of a given factor variable, we merged some levels with poor data coverage together for the *Sampling* and *Quality* variables. For instance, the levels “continuous” and “irregular” were merged together with the level “multiple” such that *Sampling* was used in our analyses as a factor variable with two levels: “two” *vs*. “multiple”. Regarding the *Quality* variable, we merged the level “resurveyed” together with the level “balanced” such that *Quality* was used in our analyses as a factor variable with three levels: “low”; “balanced”; and “modeled”.

### Taxonomic harmonization

Before undertaking any taxonomic harmonization procedure, the last version of our database, dated April 2018, contained 13,570 entries of taxa at any taxonomic rank up to the genus level (i.e. subspecies, species and genus). Using the R programming language (*23*), we assembled an R script in order to retrieve, for each taxonomic entry, the most recent accepted name and its associated classification. After a visual inspection for obvious syntax correction, three steps of taxonomic verification were performed. First, names were searched in the National Center for Biotechnology Information (NCBI) taxonomy database using the function “classification” from the R package “taxize” (*24*). Then, in the same way, any remaining taxonomic entity not found in NCBI was verified with the Integrated Taxonomic Information System (ITIS) database. The full taxonomic classification was also retrieved during these two steps. Third, the last remaining taxonomic entities not found in NCBI and ITIS were checked using the Global Biodiversity Information Facility (GBIF) database, using the function “name_backbone” in the R package “rgbif”. If we found a match, the corrected taxonomic entity was again checked in NCBI and ITIS by undergoing the previously mentioned procedure once again to retrieve a reliable taxonomic classification. Finally, only names at the species and the genus level were kept for the analyses (subspecies being aggregated at the species level). Following this taxonomic harmonization procedure, the final number of taxa names in the database was reduced to 12,415.

### Climate velocity

Using the spatial information obtained from the digitized polygons as well as the temporal information (*Start* and *End* years) available from each of the 258 publication sources (Table S1), we retrieved basic temperature information to calculate the velocity of temperature change throughout the study period. Terrestrial climate data were obtained from WorldClim v. 1.4 (http://www.worldclim.org/) and the Climate Research Unit (CRU) TS v. 3.23 (https://crudata.uea.ac.uk/cru/data/hrg/) while marine climate data were obtained from BIO-ORACLE (http://www.bio-oracle.org/) and the Met office Hadley Centre observations datasets (https://www.metoffice.gov.uk/hadobs/hadisst/).

Because marine and terrestrial taxa shift at different rates and directions to potentially track the complex mosaic of local climate velocities (*8*), we calculated the observed local velocity of temperature change (i.e. the spatial shift of isotherms over time) (*6, 7*) for each case study, following the approach used by Burrows *et al*. (*7*). We divided the temporal change in annual mean temperature observed over the studied period (°C.yr^−1^) by the corresponding spatial gradient (°C.km^−1^ or °C.m^−1^) as a measure of the velocity of temperature change (km.yr^−1^ or m.yr^−1^) (*6*). The temporal gradient was calculated using time-series data from the CRU covering the period 1901-2016 at a spatial resolution of 0.5° (about 55 km at the equator) and from the Met office Hadley Centre observations datasets covering the period 1870-2018 at a spatial resolution of 1° (about 111 km at the equator) for terrestrial and marine studies, respectively. To do so, we regressed annual mean temperature (°C) values for all years throughout the study period as well as the two preceding years against time (yr) using linear regressions. When the starting year was prior to 1901 or 1870 for terrestrial and marine systems, respectively, we started the time series in 1901 or 1870 depending on the climate series. The slope parameter (°C.yr^−1^) of this model was then used as an estimate of the temporal gradient. For the sake of comparison with the rate of range shift usually calculated along the latitudinal and elevational gradients, we calculated the spatial gradient of annual mean temperature along the latitudinal (km.yr^−1^) and elevational (m.yr^−1^) gradient, separately. This allowed us to assess both the latitudinal and elevational velocity of temperature change (*LatVeloT* and *EleVeloT*). To assess the latitudinal spatial gradient of annual mean temperature across land and sea, we used spatial grids from WorldClim and BIO-ORACLE, respectively, at a spatial resolution of 5 arc-minute (about 9.2 km at the equator). The WorldClim grid of annual mean temperature was downloaded at the finest spatial resolution, which is 30 arc-second (about 1 km at the equator), but aggregated at 5 arc-minute to be consistent with the spatial resolution of sea surface temperatures. Latitudinal spatial gradients were calculated as in Burrows *et al*. (*7*) based on a 3 × 3 neighborhood sub-grid with the centre cell being the focal cell and its eight neighboring cells used to calculate the difference in temperatures for each northern and southern (resp. southern and northern in the southern hemisphere) pairs divided by the distance between them. Average differences (°C.km^−1^) for the focal centre cell were calculated, excluding any missing values (usually along coastlines), using weightings of 1 and 2 for cells diagonal and adjacent, respectively, to the focal centre cell. For the elevation gradient, we used the temperature data from the WorldClim grid of annual mean temperature at the finest spatial resolution (30 arc-second which is about 1 km at the equator) and calculated the spatial gradient across each case study using a linear model relating annual mean temperature (the response variable) to both elevation and latitude (the explanatory variables). We used latitude as a covariate in this model to account for the latitudinal variation in temperature observed within studies covering large spatial extents, i.e. elevation values close to the equator are not directly comparable, in terms of temperature, to elevation values close to the poles. The coefficient parameter along the elevational gradient (°C.m^−1^) was then used as an estimate of the local adiabatic lapse rate. For the study areas that were larger in extent than the spatial resolution of the temperature grids, we computed the mean values of *LatVeloT* or *EleVeloT* throughout the entire study area by averaging values across all spatial grid cells overlapping with the polygons delineating the study area.

### Additional drivers of range shifts

As baseline temperature conditions may affect the rate at which species are shifting their distributions (*25*) and the rate at which communities are reshuffled (*19*), we extracted annual mean temperature values during the year of the initial census (*Start*) as well as the two preceding years and calculated the mean (hereafter *BaselineT* in °C). For terrestrial and marine systems, we used time-series data from CRU and the Met office Hadley Centre observations datasets, respectively. When the initial census of a given publication source was prior to 1901 or 1870 for terrestrial and marine systems, respectively, we used the oldest years available from the time series to compute baseline temperature conditions. Similar to climate velocity variables, when the study areas were larger in extent than the spatial resolution of the temperature grids, we computed the mean values of *BaselineT* throughout the entire study area by averaging values across all spatial grid cells overlapping with the polygons delineating the study area.

As anthropogenic disturbances such as land-use intensity or industrial fishing may act as confounding factors on the rate of range shift (*25*), we retrieved information on anthropogenic impacts for both the terrestrial and marine environment. For terrestrial systems, we downloaded the Global terrestrial Human Footprint maps for the year 2009 (*13*). These maps, at a spatial resolution of 30 arc-second (about 1 km at the equator), provide remotely-sensed and bottom-up survey information on eight variables measuring the direct and indirect human pressures on the environment: (1) built environments; (2) population density;(3) electric infrastructure; (4) crop lands; (5) pasture lands; (6) roads; (7) railways; and navigable waterways. All eight pressure maps were cumulated into a global human footprint index ranging from 0 to 50. For marine systems, we used the Global Map of Human Impact on Marine Ecosystems (*12*), also available at 30 arc-second resolution (about 1 km at the equator). This gridded dataset provides a cumulative impact score ranging from 0.01 to 90.1 for the minimum and maximum value, respectively. It was developed on the basis of expert judgment, to estimate ecosystem-specific impacts with respect to 17 anthropogenic drivers of ecological change (e.g. commercial shipping, demersal and pelagic fishing, ocean acidification, pollution). To allow comparison between terrestrial and marine systems, we rescaled both indices between 0 and 1 (standardized human footprint index or standardized *HFI*) and computed the mean per study area.

### Description: assessing geographic and taxonomic biases

To evaluate spatial biases in the reporting of range shift rates, we built 2° × 2° gridded maps, on top of which we overlaid the digitized polygons associated with the observations gathered in the database for both the terrestrial and marine realm, and separately for latitudinal and elevational range shifts. For each 2° × 2° grid cell, we also computed the relative proportion of ectotherms *vs*. endotherms for animals and phanerogams *vs*. cryptogams for plants and plant-like life forms (e.g. lichens and algae). We distinguished ectotherm from endotherm life-forms due to their contrasting sensitivity to temperature fluctuations in the environment, with ectotherms being unable to directly regulate their body temperatures as opposed to endotherms. The distinction between phanerogams and cryptogam life-forms allowed to contrast between two different reproduction strategies among chlorophyllous organisms: the evolved seed-bearing plants (angiosperms and gymnosperms) *vs*. the other plant-like life forms reproducing by spores (ferns, mosses, lichens and algae). We then generated cartograms using the diffusion-based method for producing density-equalizing maps (*26*). The number of range shift rates per 2° × 2° grid cell (i.e. sample size) was used to distort the map: the bigger the grid cell, the larger the sample size (see Fig. S1). We additionally estimated the phylogenetic coverage of the range shift database with respect to the whole tree of life described in the Open Tree of Life (https://tree.opentreeoflife.org) collapsed at the level of taxonomic classes and the total number of species recorded in the Catalogue of Life (http://catalogueoflife.org/).

### Detection: estimating the velocity of range shifts per taxonomic class

Data coverage in our database is very unbalanced between: the marine *vs*. terrestrial realm; the northern *vs*. southern hemisphere; and the margins *vs*. centroid of the species range (Table S2). Besides, data on species range shifts do not even exist for some taxonomic classes in some of the combination of realm × hemisphere × position in the species range. For instance, dicots (*Magnoliopsida*) are exclusively terrestrial organisms while cartilaginous fishes (*Chondrichthyes*) almost exclusively live in marine habitats except for a few sharks and rays living in freshwater habitats during all or part of their lives. Thus, a single model to estimate the velocity of range shifts per taxonomic class while accounting for methodological biases (*4, 11, 22*) would be inappropriate. Hence, we divided latitudinal range shifts (N = 16,952) into a full factorial design (*27*) with eight experimental units based on all possible combinations of levels across three factor variables: biological system (marine *vs*. terrestrial); hemisphere (north *vs*. south); and range position (centroid *vs*. margins). We did the same for elevational range shifts (N = 13,582) except that there were only four possible experimental units (i.e. terrestrial systems only). To ensure robust fit, we further focused on taxonomic classes with more than 30 observations per experimental unit (N = 20 taxonomic classes fulfilling this sample size criterion; Table S2; Fig. 1), which reduced our sample size to 16,399 and 13,341 observations for latitudinal and elevational range shifts, respectively. Among the 12 possible combinations, only one combination (latitude × margins × terrestrial × south) could not be fulfilled due to a lack of range shift data (N = 8). This resulted in a total of 11 sub-models (i.e. factorial models) (Table S2).

For each of the 11 factorial models for which the data were available, we built a linear mixed-effects model (LMM) relating the velocity of species range shift (*ShiftR*) for a given taxon (i.e. the response variable) against taxonomic *Class*, a factor variable with as many levels as is the number of taxonomic classes within the focal experimental unit (e.g. *Amphibia vs. Aves* for latitudinal range shifts at the centroid of the distribution in terrestrial systems of the southern hemisphere) (Table S2). Note that if a given factorial model only had data for one unique taxonomic class (e.g. *Actinopterygii* for latitudinal range shifts at the centroid of the distribution in marine systems of the southern hemisphere) (Table S2), then the variable *Class* was not included in the fixed effects of the LMM. For the five LMMs focusing on the rate of range shift at the margins of the distribution, we added an extra factor variable (*Margin*) with two levels (“leading” *vs*. “trailing” edge) in the fixed effects, to provide robust estimates of the rate of range shift at both the leading and trailing edges. Given the complex structure of the database, sometimes involving several observations for a given family or genus or per level of a methodological variable, LMM is the most appropriate modelling approach (*27*). This allowed to provide estimates of the velocity of range shifts per taxonomic class that are representative across all levels of a given methodological variable rather than providing estimates for each level separately. More specifically, we included *Genus* as a random intercept term nested (or not: in case of singularity fit) within *Family* to account for potential taxonomic autocorrelation in the residuals of the models. In addition, because the different methodological approaches used in the scientific studies may also contribute to a non-negligible fraction of the variation in range shifts (*11*), we allowed several methodological variables to be included as random intercept terms in the LMMs (*Area, Start, Ntaxa, Sampling, PrAb, Grain, Quality* and *Signif*). To be potentially included in the random part of our LMMs, the continuous variables *Area, Start* and *Ntaxa* were first transformed into factor variables with four levels each, using the respective quantiles as cutting points. Then, for each factorial model separately, we selected only the set of uncorrelated variables with at least two levels having more than four observations. We used the “lmer” function from the “lme4” package (*28*) in the R programming language (*23*).

Regarding model selection strategy, the best random effect structure was identified by testing every variable combinations and selecting the one with the lowest Akaike information criterion (AIC) value, with small-sample correction (AICc). To compare AICc values among candidate models, we set the restricted maximum likelihood argument to “FALSE” in the “lmer” function (i.e. REML = FALSE for maximum likelihood) (*28*). To ensure robust estimations, all the singularity fits were removed from the list of candidate models prior to model selection. In case of singularity fits across all candidate models testing all combinations of variables in the random effect structure, we used case study (*Source*) as the unique random intercept term. Finally, if the random intercept term *Source* also led to a singularity fit, then we used a simple linear regression model (LM). For each of the LMMs (or LMs in case of singularity fits for all tested random effect structures) focusing on the rate of range shift at the margins of the distribution, we then tested the interaction effect between *Margin* and *Class* against the model without the interaction term in the fixed effects. Again, the best fixed effect structure for these two candidate models was selected based on the AICc value. When the absolute difference in AICc value between these two candidate models was greater than two, we selected the model with the lowest AICc value. Otherwise, in case of equivalent competing models, we selected the one with the interaction effect between *Margin* and *Class* considering that it allows flexible range shift estimations at the trailing and leading edge. Once the best LMM was selected for each factorial model (see Table S2), we set REML to TRUE (*28*) to extract coefficient estimates among the different levels of the factor variables *Class* and *Margin*. To test whether the estimated rate of range shift for a given taxonomic class and at a given position within the range was significantly different from zero, we reran each of the 11 selected best models using a bootstrap approach (N = 5,000 iterations). From these 5,000 estimates, we computed the mean and median velocity of range shift as well as the standard deviation and 95% confidence interval per taxonomic class. Finally, to assess the overall goodness-of-fit of the different factorial models, as well as to compare the relative importance of biological versus methodological effects on the rate of range shift, we computed the marginal (i.e. variance explained by the fixed effects) and conditional (i.e. variance explained by both the fixed and random effects) R^2^ (*29*) of each bootstrap iteration and for each factorial model using the “r.squaredGLMM” function from the “MuMIn” package in the R programming language (*23*).

### Attribution: coupling between species’ range shifts and isotherms’ shifts

We assessed the coupling between the velocity of species range shifts and the velocity of isotherm shifts using LMMs built separately for the latitudinal and the elevational gradients. We specified the velocity of species latitudinal (km.yr^−1^) or elevational (m.yr^−1^) range shifts as the response variable and either the latitudinal velocity of isotherm shifts or the elevational velocity of isotherm shifts (*LatVeloT* / *EleVeloT*; continuous variables) as the main predictor variable. To account for potential interacting effects on the relationship between the velocity of range shifts and the velocity of isotherm shifts, we added several covariates in our models: baseline temperature (*BaselineT*; a continuous variable); human footprint index (standardized *HFI*; a continuous variable representing human pressures on the environment bounded between 0 and 1); and *LifeForm* (a factor variable with 4 levels: ectotherm, endotherm, cryptogam, phanerogam). As temperature regimes and human pressures on the environment are not directly comparable between lands and oceans, we further modeled the coupling between the velocity of species latitudinal range shifts and the velocity of isotherm shifts in latitude separately for the marine and terrestrial realm. We tested for all two-way interaction terms between each covariate (*BaselineT, HFI* and *LifeForm*) and the velocity of isotherm shifts, either *LatVeloT* or *EleVeloT*. We also tested for a unimodal relationship between the estimated rates of range shifts and baseline temperature conditions (*BaselineT*) using a second-order polynomial term. The variables *Position* within the range (a factor variable with 3 levels: trailing edge, centroid and leading edge) and *Hemisphere* (a factor variable with 2 levels: North *vs*. South) were not incorporated as covariates in the models as both variables had no effect to explain the variation in the rates of latitudinal and elevational range shifts per taxonomic class.

Similar to the LMMs developed at the taxonomic class, the aforementioned predictors were used as fixed effects in LMMs, whereas the methodological attributes (*Area, Start, Ntaxa, Sampling, PrAb, Grain, Quality* and *Signif*) were used as random intercept terms. Starting from the beyond optimal model (full model with all fixed effects) (*28*) for the velocity of latitudinal range shifts in marine and terrestrial systems as well as for the velocity of elevational range shifts, we tested all model combinations and selected the best model based on the lowest AICc value, setting REML to FALSE (i.e. maximum likelihood) for model selection and then to TRUE for coefficient estimates once the best model was selected (*28*). We first selected the random effect structure after removing singularity fits, using the exact same procedure as for the models used to estimate the mean velocity of range shift per taxonomic class, and then the fixed effect structure, keeping the previously identified random structure constant. All continuous variables (*LatVeloT, EleVeloT, BaselineT* and *HFI*) were standardized to *z*-scores using the “gscale” function (*30*) from the “jtools” package in the R programming language (*23*). This function standardizes each value of a given variable by subtracting it from the mean and dividing it by two times, instead of one time, the standard deviation of the focal variable. This rescaling formula is recommended over the traditional formula of dividing by one time the standard deviation because it allows direct comparisons of model coefficients with untransformed binary predictors (*30*). For the sake of consistency, we focused on the set of species belonging to the taxonomic classes with more than 30 observations, resulting in 16,521 (1,403 marine *vs*. 15,118 terrestrial) and 13,459 observations for latitudinal and elevational range shifts, respectively. The 95% confidence intervals around each of the estimated coefficients were calculated through a bootstrap approach (N = 5,000 iterations), using the exact same approach as for the models used to estimate the mean velocity of range shift per taxonomic class.

Finally, to illustrate the species’ ability to closely track the shifting isotherms, we mapped the predicted slopes for each combination of the predictors identified in the best models, separately for latitudinal (marine or terrestrial) and elevational range shifts. A slope value of one between the velocity of species range shifts and the velocity of isotherm shifts indicates a perfect coupling with species closely tracking their shifting isotherms. To do so, we built a 2° × 2° gridded map, on top of which we overlaid the digitized polygons associated with each observation used in the previous models. We then generated cartograms using the diffusion-based method for producing density-equalizing maps (*26*). As before, the number of range shift rates per grid cell (i.e. sample size) was used to distort the map: the bigger the grid cell, the larger the sample size. For each 2° × 2° grid cell, we then tested whether the predicted slope parameter, based on the information available within the focal grid cell (i.e. baseline temperature and the standardized human footprint index), did significantly differ or not from a value of one (perfect coupling), based on 5,000 bootstrap iterations.

## Acknowledgments

We acknowledge authors who kindly sent us their data on range shifts estimates; We acknowledge Investissement d’Avenir grants of the Agence Nationale de la Recherche (TULIP ANR-10-LABX-41, CEBA ANR-10-LABX-25-01); Author contributions: JL, LC, JM and GG initiated and conceived the project idea. LC and JL built the general structure of the database. GG, LC, RB, TH and JL reviewed the scientific literature and filled the database throughout the project duration. GG ensured data curation. LB and LC carried out the taxonomic harmonization of the database with help from JM. TH linked the taxonomic backbone of the database to the Open Tree of Life (https://tree.opentreeoflife.org) and to Catalogue of Life (http://catalogueoflife.org/) to produce a visualization of the phylogenetic coverage of the database. GG, LC, JL and RB prepared the set of methodological variables included as covariates in the subsequent analyses. RB and JL analyzed the data with help from LC, LB and GG. TH, RB and JL produced all the figures. JL wrote the manuscript with contribution from all co-authors. JL and RB contributed equally. Competing interests: Authors declare no competing interests; and Data and materials availability: The entire range-shift database and R scripts used in the analyses will be made available online, as soon as the manuscript will be accepted for publication.

## Supplementary Discussion

### Geographical bias in our knowledge of climate-induced range shifts

For latitudinal range shifts on lands and in the oceans, our analysis confirms an important knowledge gap in the tropics (*4, 23*). Indeed, there is a strong spatial unbalance in the database, such that the most developed regions of the northern hemisphere are better represented in terms of latitudinal range shifts: Western Europe; North America; the North Atlantic Ocean; and the North Sea (Fig. S1). By contrast, the southern hemisphere is underrepresented with Australia and the Tasmanian Sea being the most studied regions. We observed a similar pattern for elevational range shifts on lands. Again, mountain ranges from the most developed countries are better covered (i.e. Rockies and European Alps) than elsewhere, with tropical mountains poorly represented. Hence, the geographical coverage of the data on species range shifts is far from global (*23*). Finally, there is also a geographical bias among the different life forms (i.e. ectotherms, endotherms, phanerogams and cryptogams) covered in the database. Ectotherms are overrepresented for latitudinal range shifts in marine systems, whereas data on latitudinal range shifts for endotherms (mostly birds) dominate in North America, Finland and Australia (Fig. S2) where active groups of researchers are working on well-maintained databases of bird censuses.

### Taxonomic bias in our knowledge of climate-induced range shifts

Despite a broad taxonomic coverage of the tree of life (Fig. S3), species range shifts have only been documented for 0.6% of the described biodiversity on Earth (N = 2,094,892 taxa). There is a clear bias to report range shifts for: the most mobile ectotherms, such as flying insects (*Insecta*) and bony fishes (*Actinopterygii*); the most iconic endotherms, such as birds (*Aves*) and mammals (*Mammalia*); and seed-bearing phanerogams, such as dicots (*Magnoliopsida*) and monocots (*Liliopsida*). By contrast, cryptogams such as mushrooms (*Lecanoromycetes*) and algae (*Florideophyceae*) are poorly represented.

## Supplementary Figures

**Fig. S1.**
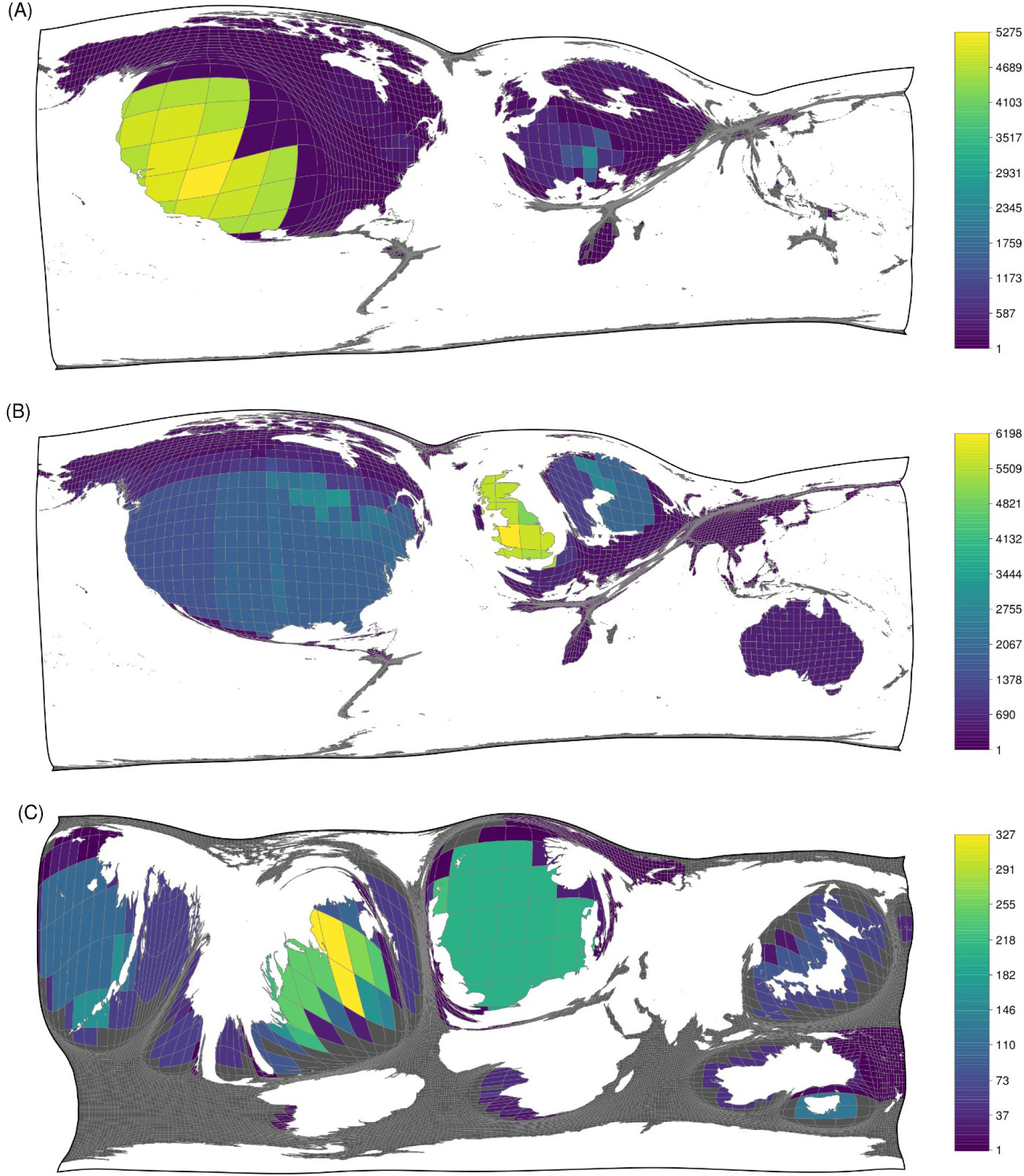
Cartograms of the number of taxa per 2° × 2° grid cell and for which we have data on (**A**) elevational range shifts and data on (**B, C**) latitudinal range shifts for both the terrestrial (**B**) and (**C**) marine realm.

**Fig. S2.**
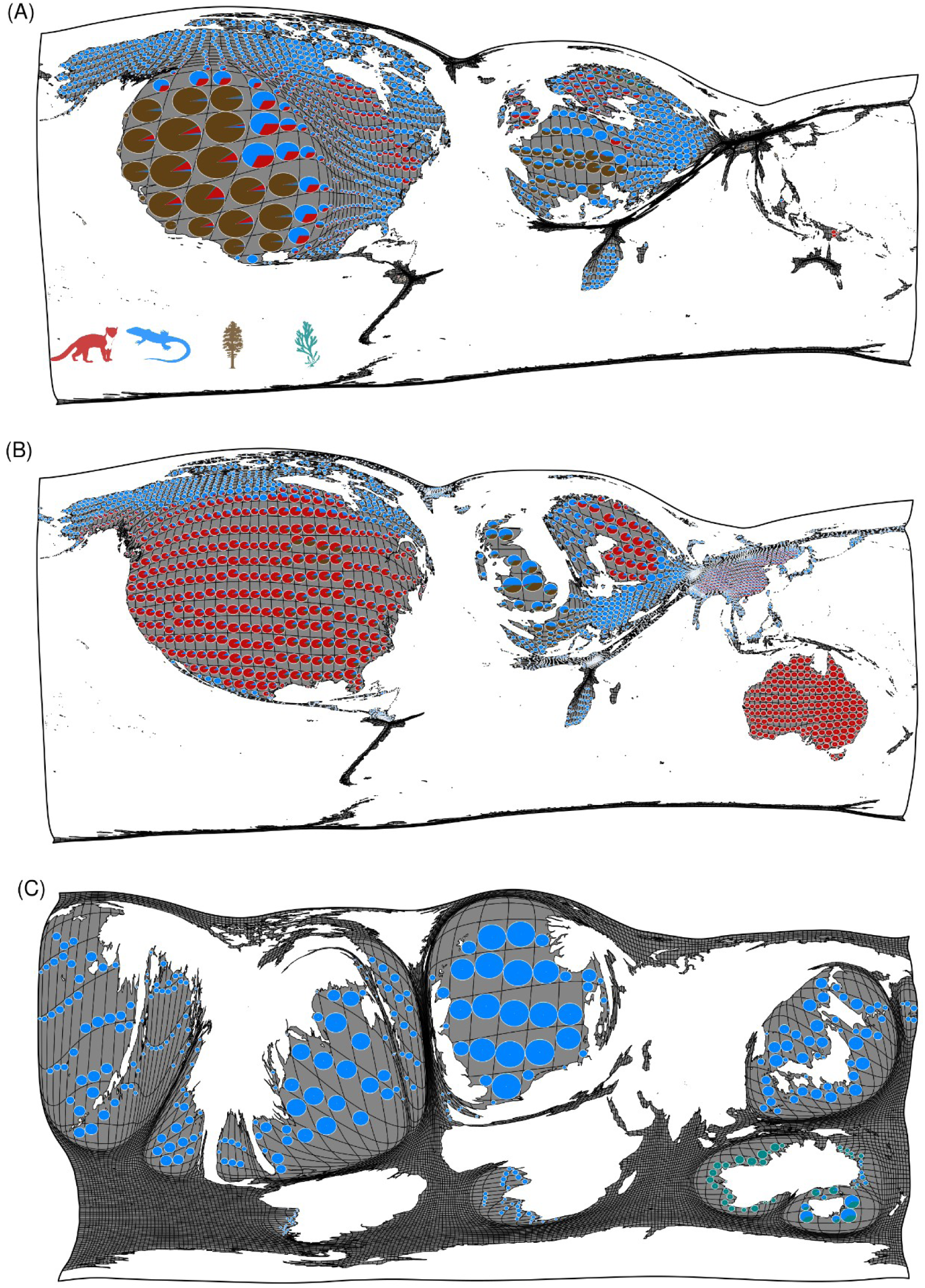
Cartograms of the relative proportion of ecotherms, endotherms, phanerogams and cryptogams per 2° × 2° grid cell and for which we have data on (**A**) elevational range shifts and data on (**B, C**) latitudinal range shifts for both the terrestrial (**B**) and (**C**) marine realm.

**Fig. S3.**
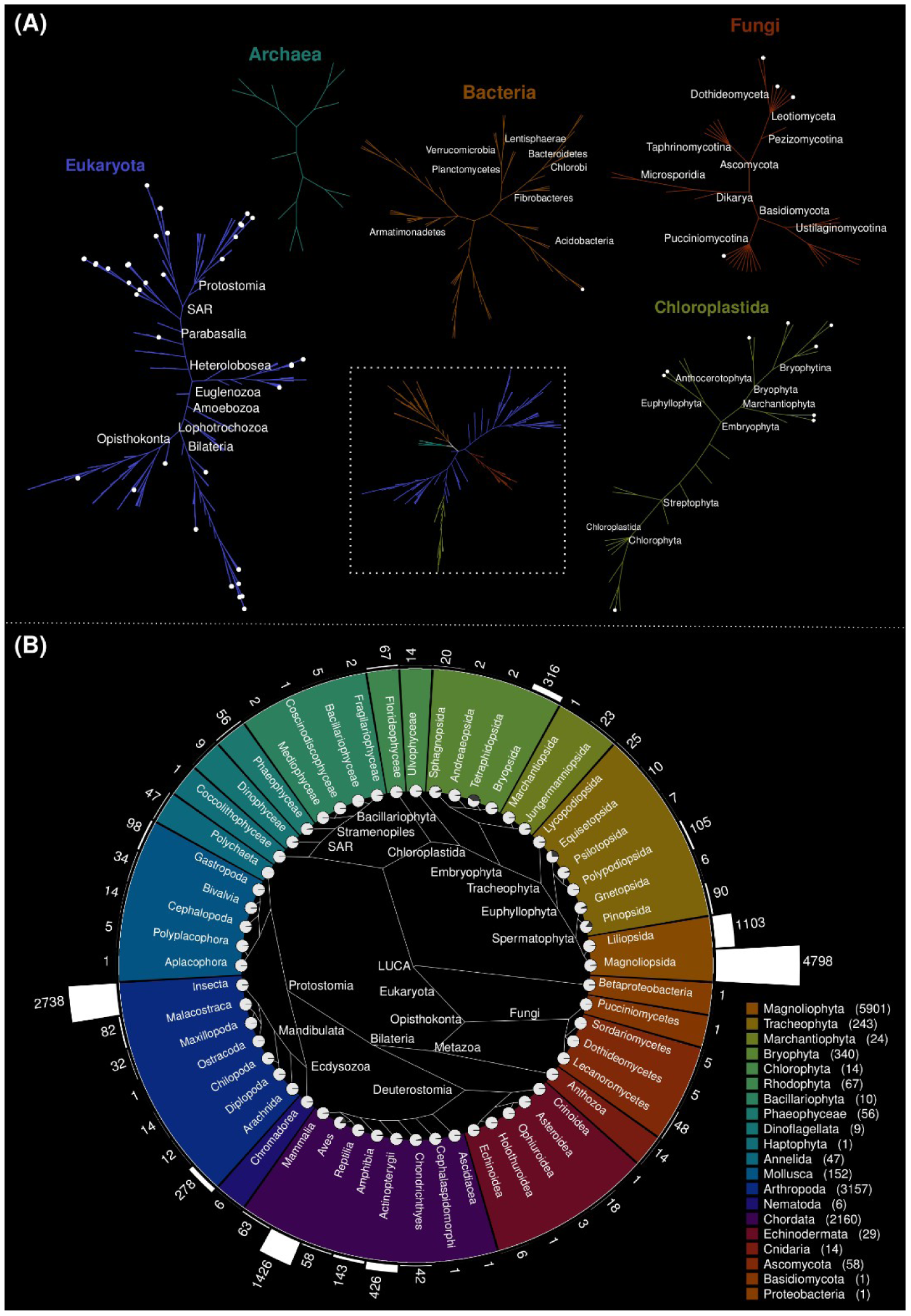
Phylogenetic coverage of the database on species range shift throughout (**A**) the whole tree of life with a focus on (**B**) the phylogenetic relationships among the 56 taxonomic classes included in the database. Simplified representation of the Open Tree of Life (https://tree.opentreeoflife.org) collapsed at the level of taxonomic classes. Clades included in the database are highlighted by white dots at the tips. Branches’ colors indicate the taxonomic phylum to which classes belong. Bars show the number of species registered in the database per taxonomic class. Pie charts at the tips of the phylogeny represent the proportion of species recorded in the database compared to the total number of species recorded in Catalogue of Life (http://catalogueoflife.org/). Colors represent the 20 phyla occurring in the database (the number of species per phyla is provided in parentheses).

**Fig. S4.**
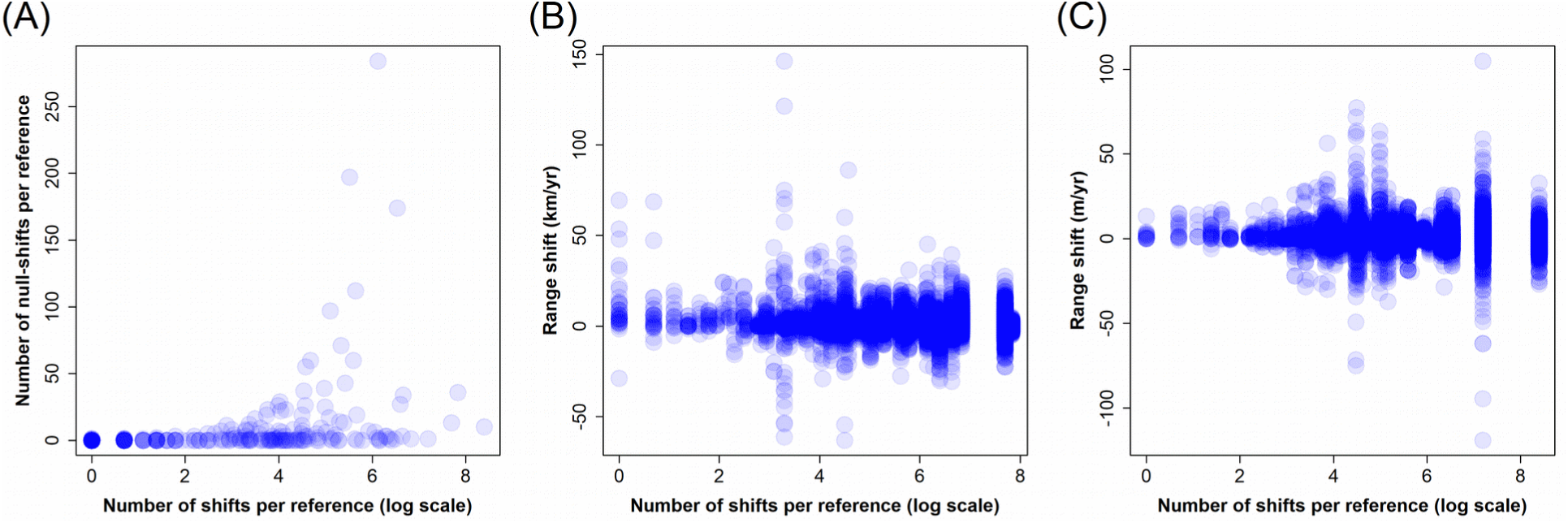
Total number of range shifts reported per study (mind the x-axis in log-scale) with (**A**) the frequency of null-shifts per study as well as its effects on the magnitude of the shift in (**B**) latitude and in (**C**) elevation.

**Fig. S5.**
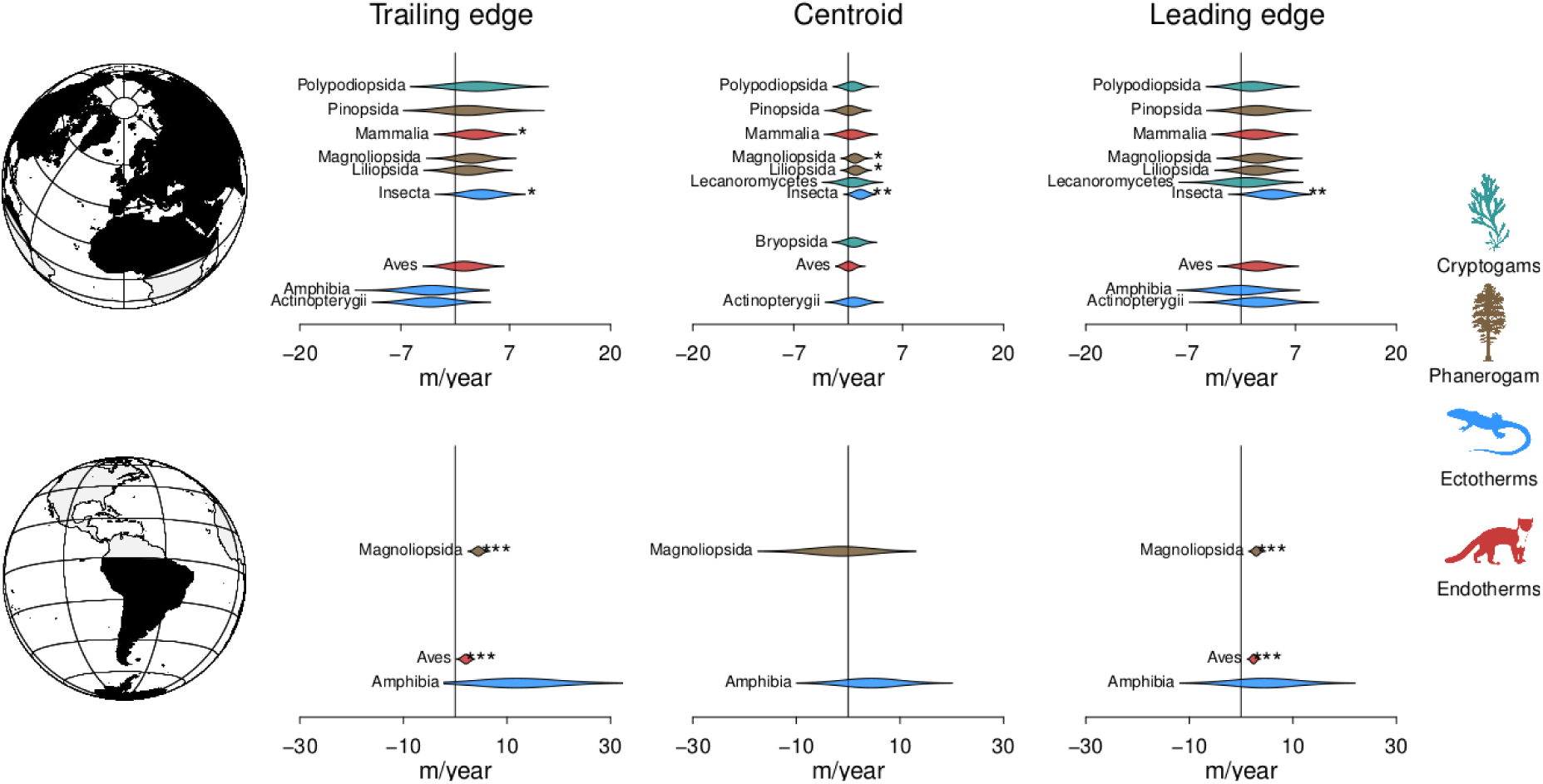
Estimated rate of elevational range shift per taxonomic class in m.yr^−1^. Violin plots represent the distribution of 5,000 bootstrap iterations. Stars show significant effects (*: *P* < 0.05; **: *P* < 0.01; ***: *P* < 0.001) determined from bootstrap distributions for the two alternative hypotheses of a mean rate of range shift being either negative or positive relative to a zero shift (i.e. the null hypothesis).

**Fig. S6.**
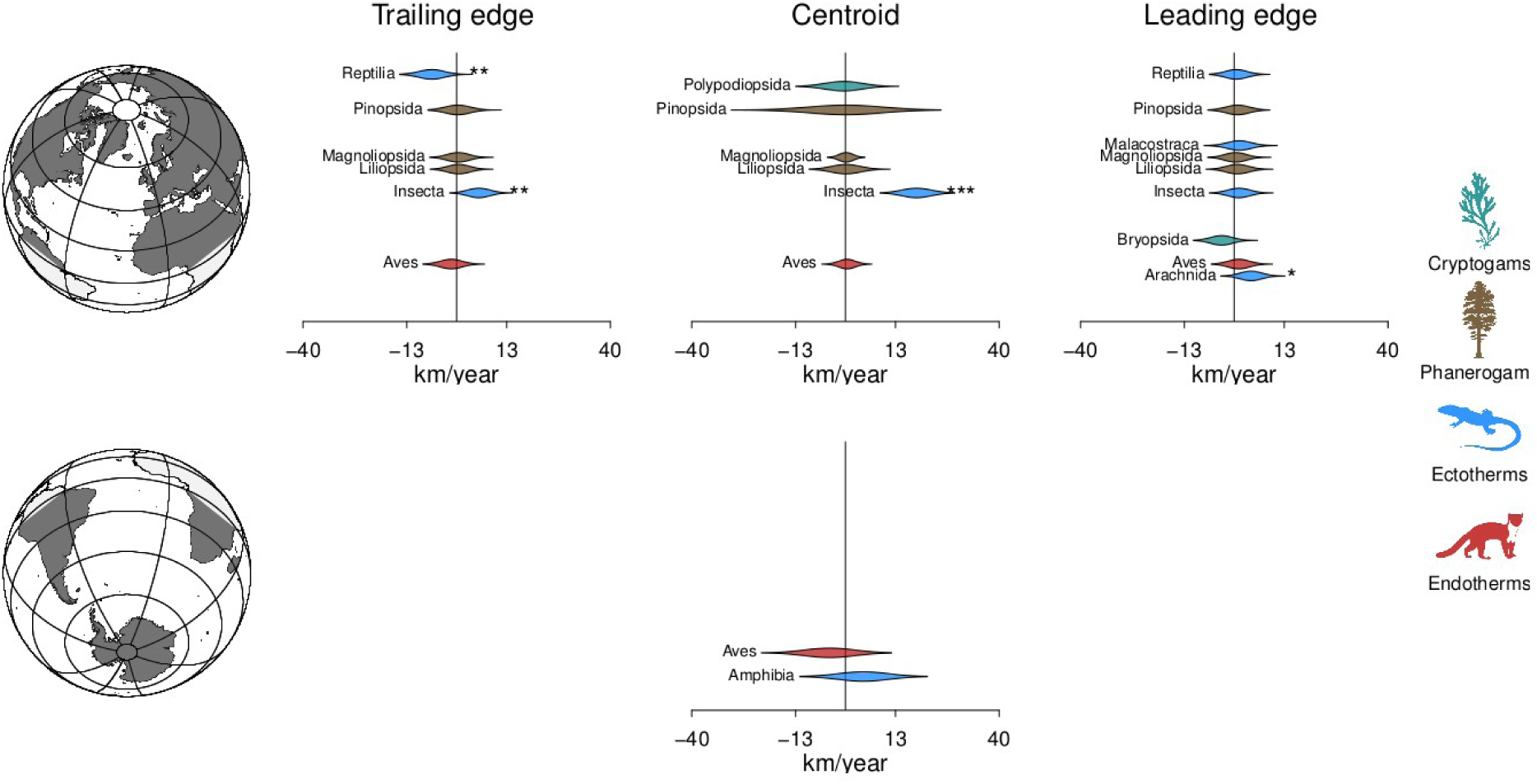
Estimated rate of latitudinal range shift per taxonomic class in km.yr^−1^ in terrestrial systems. Violin plots represent the distribution of 5,000 bootstrap iterations. Stars show significant effects (*: *P* < 0.05; **: *P* < 0.01; ***: *P* < 0.001) determined from bootstrap distributions for the two alternative hypotheses of a mean rate of range shift being either negative or positive relative to a zero shift (i.e. the null hypothesis).

**Fig. S7.**
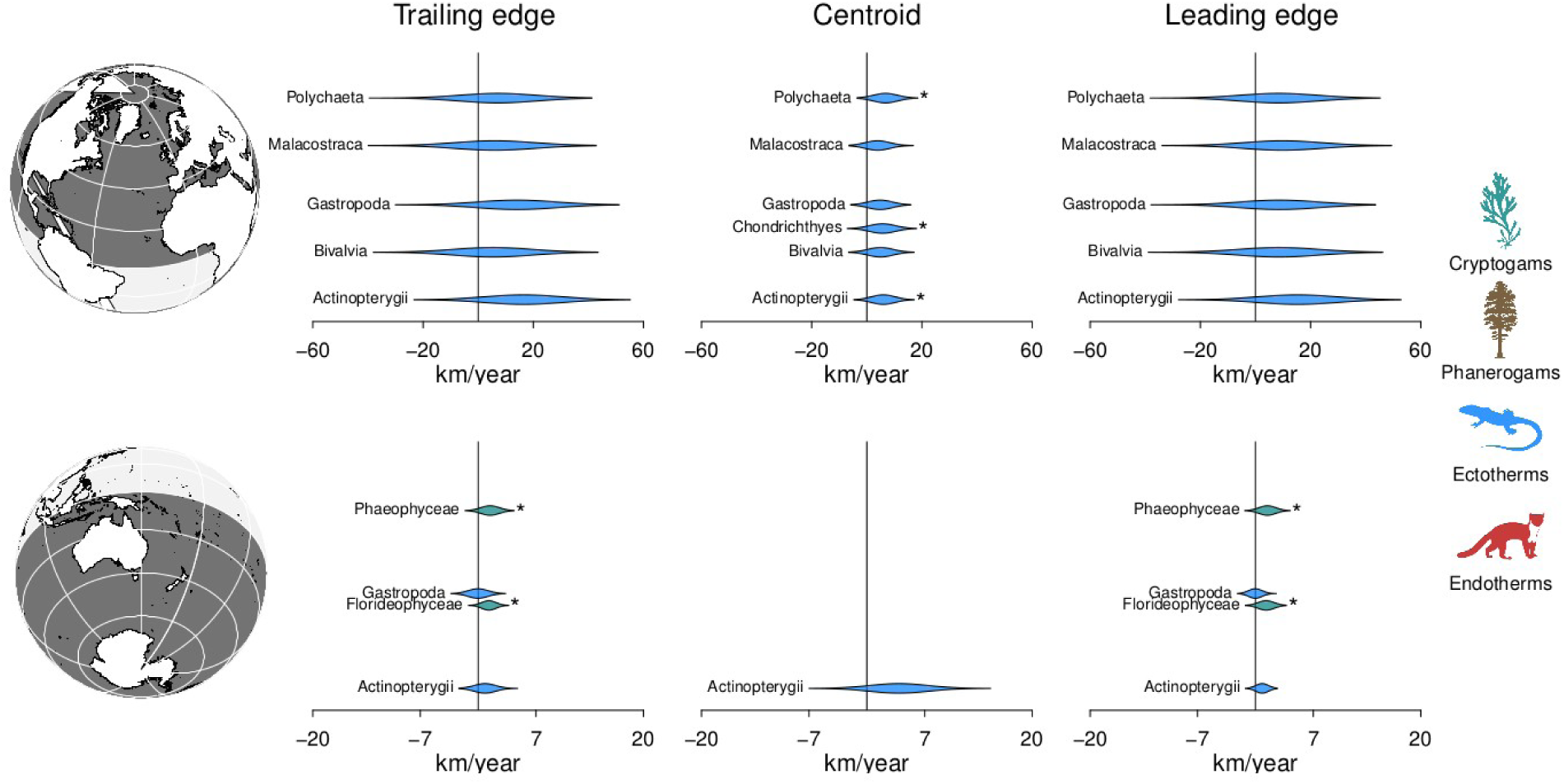
Estimated rate of latitudinal range shift per taxonomic class in km.yr^−1^ in marine systems. Violin plots represent the distribution of 5,000 bootstrap iterations. Stars show significant effects (*: *P* < 0.05; **: *P* < 0.01; ***: *P* < 0.001) determined from bootstrap distributions for the two alternative hypotheses of a mean rate of range shift being either negative or positive relative to a zero shift (i.e. the null hypothesis).

**Fig. S8.**
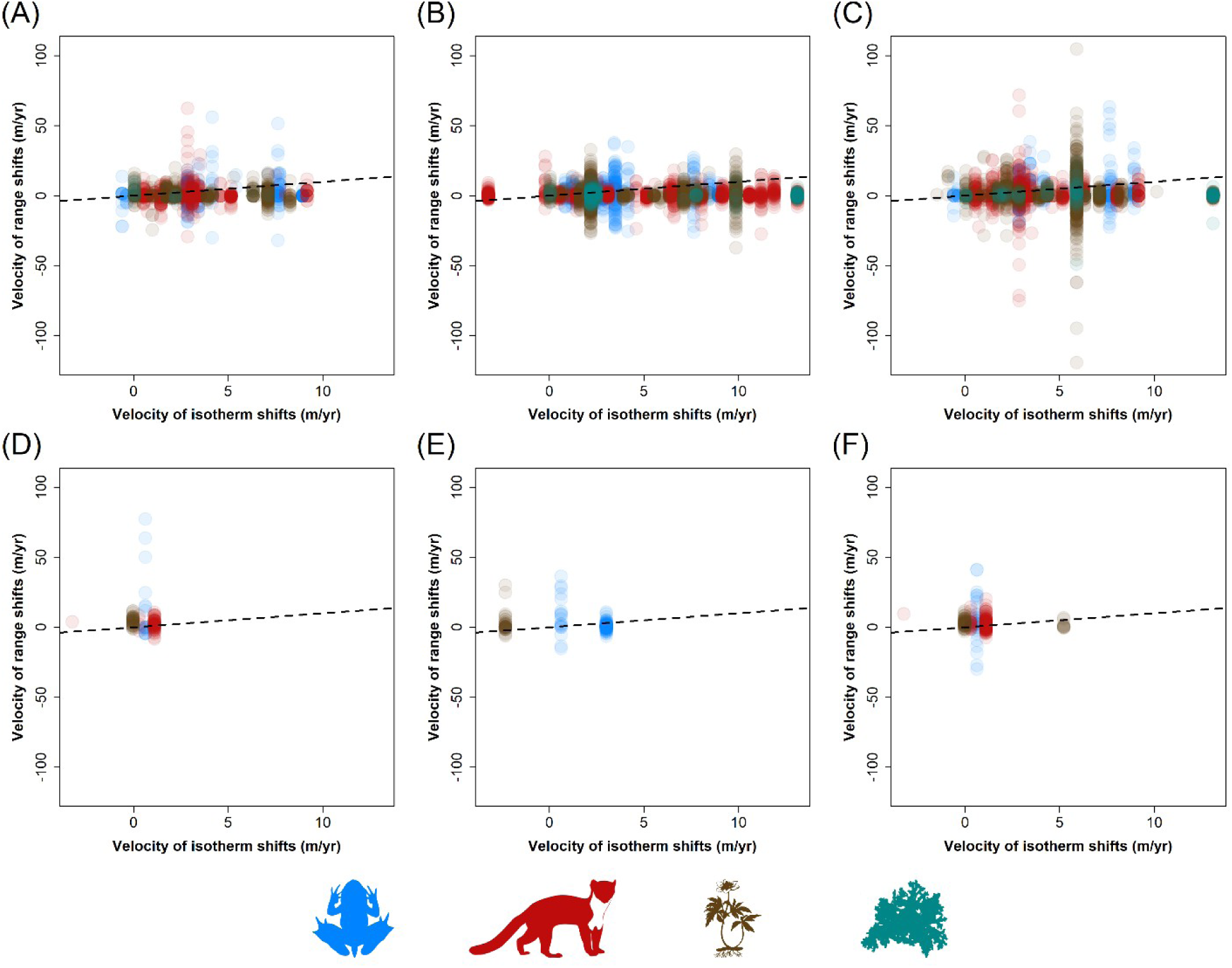
Degree of coupling between the velocity of species elevational range shifts (m.yr^−1^) the velocity of isotherms’ shifts in elevation (m.yr^−1^). The degree of coupling is displayed separately for the (**A, B, C**) northern and (**D, E, F**) southern hemisphere and separately for the (**A, D**) trailing edge, (**B, E**) centroid and (**C, F**) leading edge of the range. The dotted line represents the 1:1 relationship of perfect match, meaning that organisms are closely tracking the shifting isotherms. Ectotherms, endotherms, phanerogams and cryptogams are displayed in blue, red, brown and cyan, respectively.

**Fig. S9.**
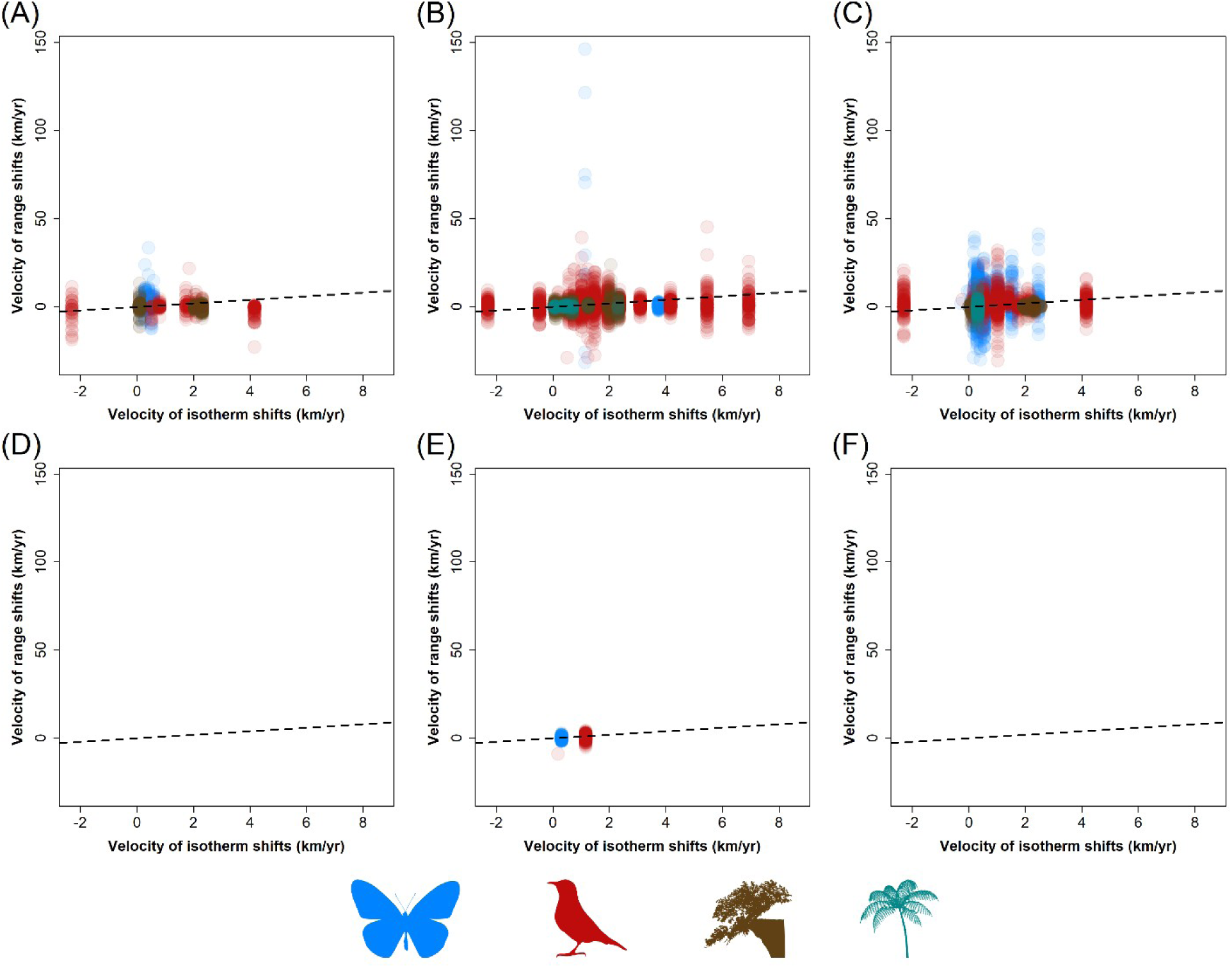
Degree of coupling between the velocity of species latitudinal range shifts (km.yr^−1^) and the velocity of isotherms’ shifts in latitude (km.yr^−1^). The degree of coupling is displayed separately for the (**A, B, C**) northern and (**D, E, F**) southern hemisphere and separately for the (**A, D**) trailing edge, (**B, E**) centroid and (**C, F**) leading edge of the range. The dotted line represents the 1:1 relationship of perfect match, meaning that organisms are closely tracking the shifting isotherms. Ectotherms, endotherms, phanerogams and cryptogams are displayed in blue, red, brown and cyan, respectively.

**Fig. S10.**
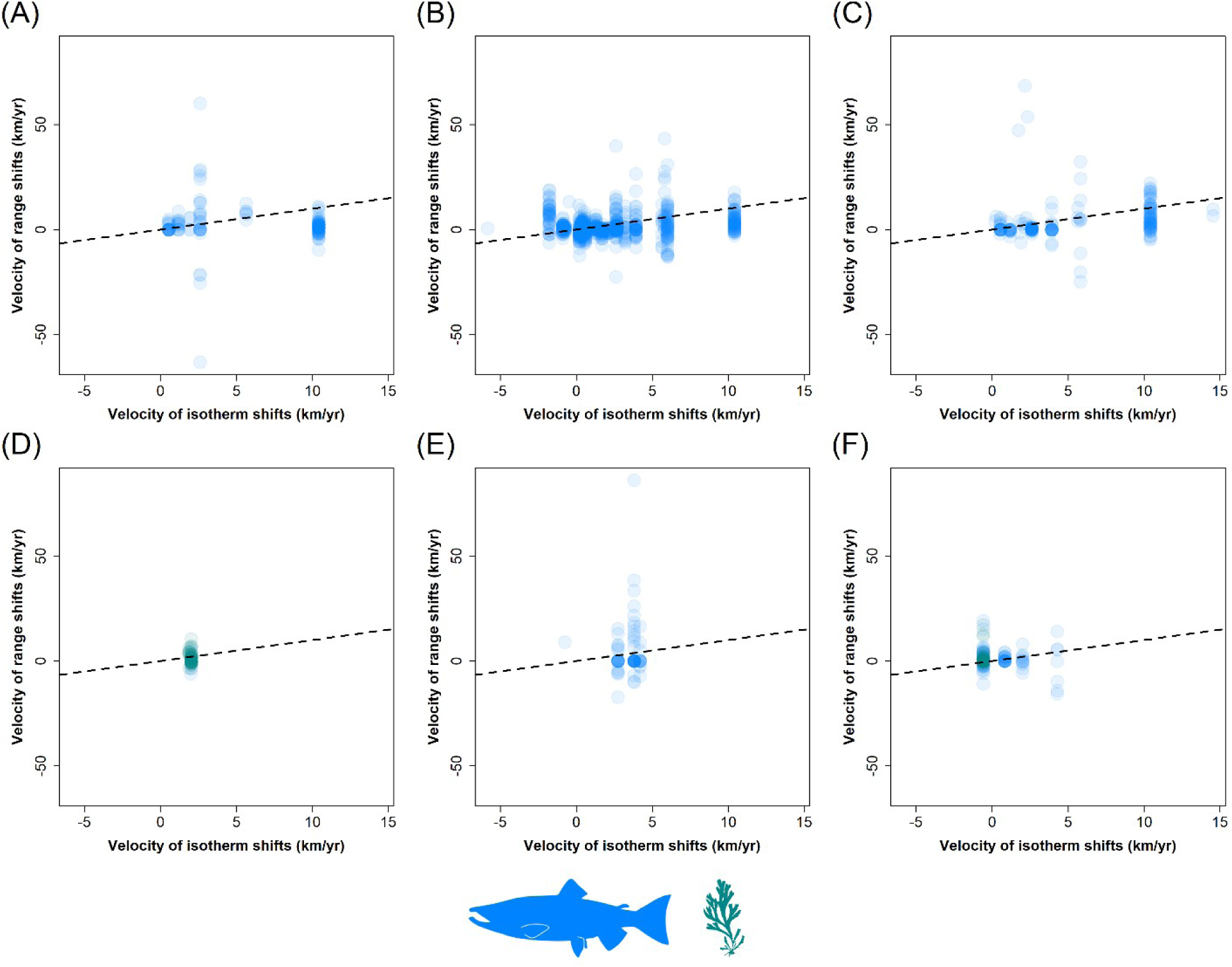
Degree of coupling between the velocity of species latitudinal range shifts (km.yr^−1^) and the velocity of isotherms’ shifts in latitude (km.yr^−1^). The degree of coupling is displayed separately for the (**A, B, C**) northern and (**D, E, F**) southern hemisphere and separately for the (**A, D**) trailing edge, (**B, E**) centroid and (**C, F**) leading edge of the range. The dotted line represents the 1:1 relationship of perfect match, meaning that organisms are closely tracking the shifting isotherms. Ectotherms and cryptogams are displayed in blue and cyan, respectively.

**Fig. S11.**
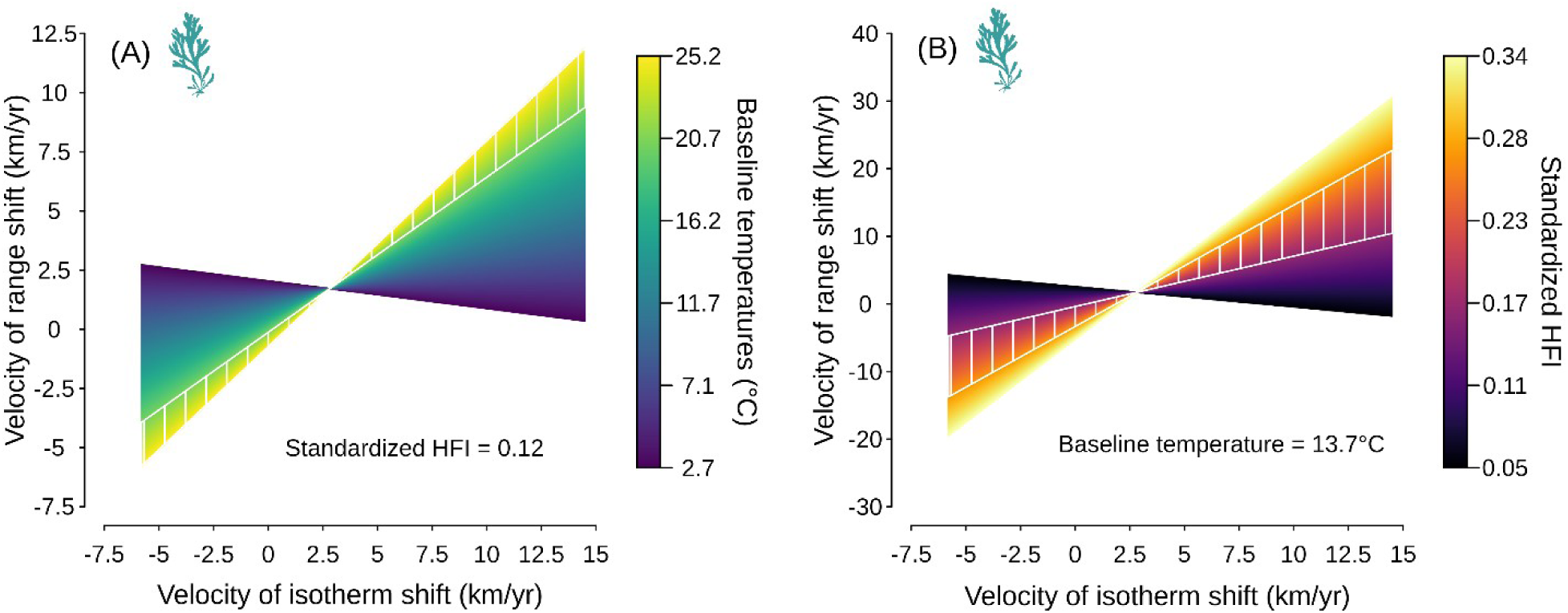
Interaction effects between the velocity of isotherm shifts and (**A**) baseline temperatures or (**B**) the standardized human footprint index on the velocity of species range shifts along the latitudinal gradients for marine cryptogams (see. Figs. 4C-4D for marine ectotherms). The two white lines and the white hatching represent the range of conditions for which marine cryptogams closely track the shifting isotherms in latitude (i.e. slope parameter not significantly different from 1 based on 5,000 bootstrap iterations).

**Fig. S12.**
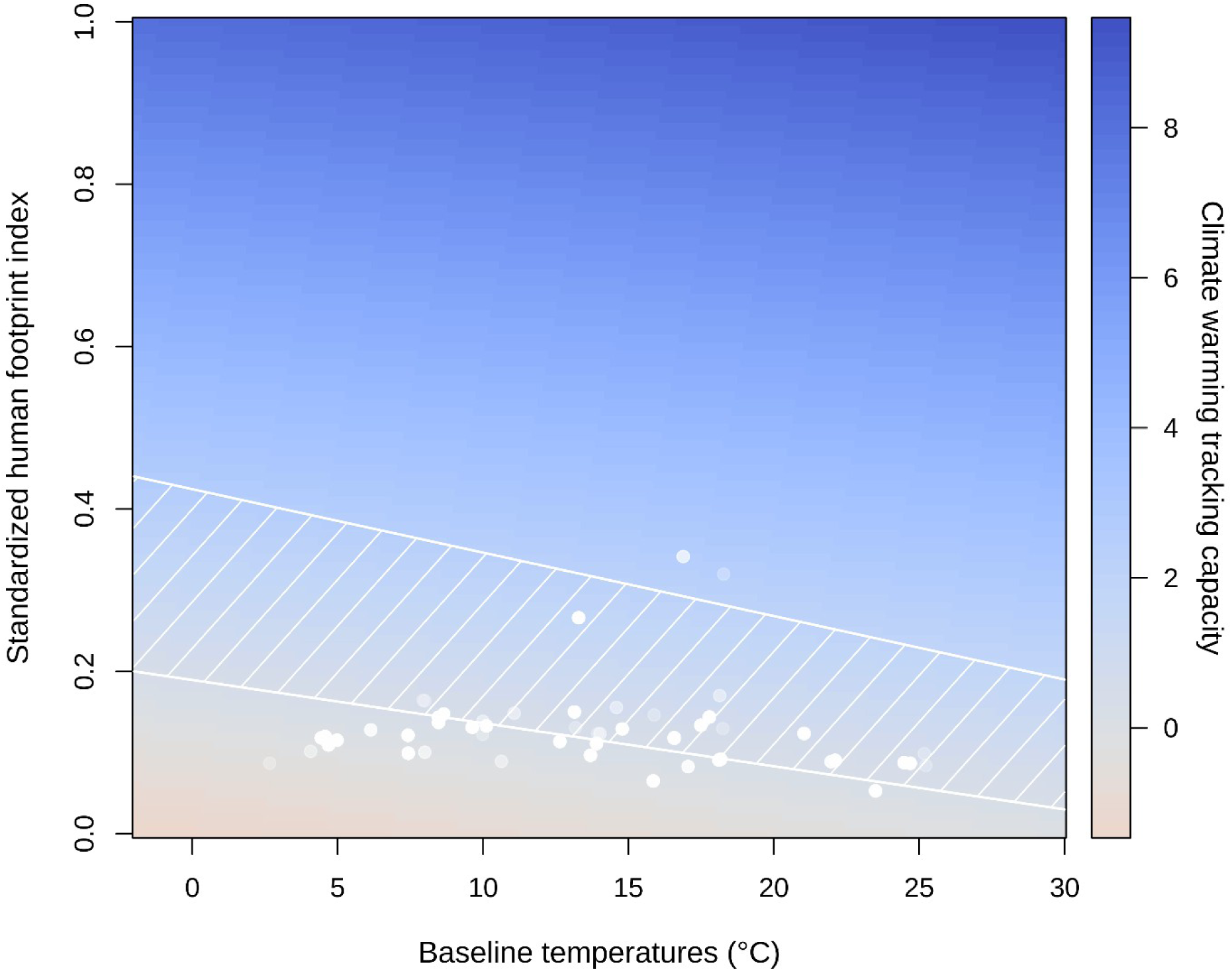
Combined effect of mean annual sea surface temperature prior to the baseline survey (baseline temperatures) and human pressures on the environment (the standardized human footprint index) on the slope of the relationship between the velocity of marine species range shifts and the velocity of isotherm shifts along the latitudinal gradient in the oceans. The white lines and hatching display the range of conditions for which the predicted slope of isotherm tracking is not significantly differing from one, based on 5,000 bootstrap iterations, meaning that marine taxa are closely tracking the shifting isotherms. White transparent dots show the distribution of the raw data (N = 1403 range shifts for marine taxa) used to fit the model. This plot applies for both marine ectotherms and cryptogams.

**Fig. S13.**
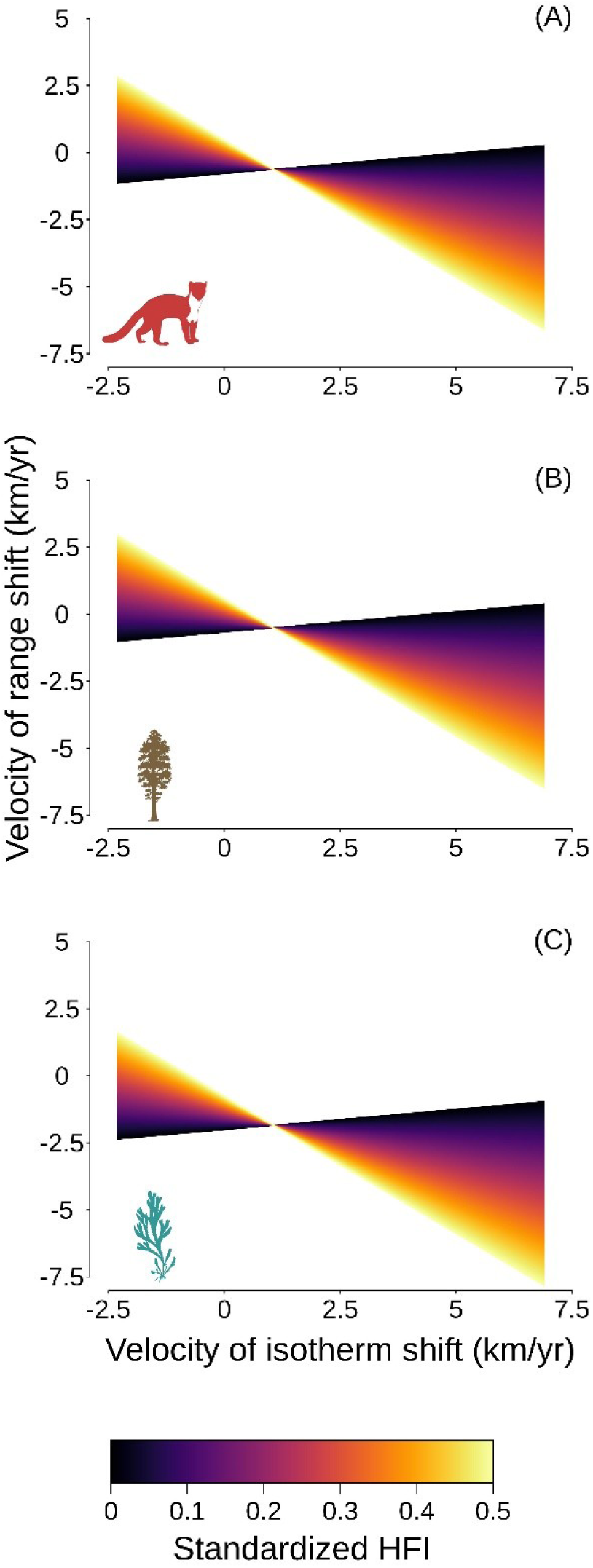
Interaction between the standardized human footprint index (HFI) and the velocity of isotherm shifts in latitude for terrestrial (**A**) endotherms, (**B**) phanerogams and (**C**) cryptogams (see Fig. 3B for the case of ectotherms).

**Fig. S14.**
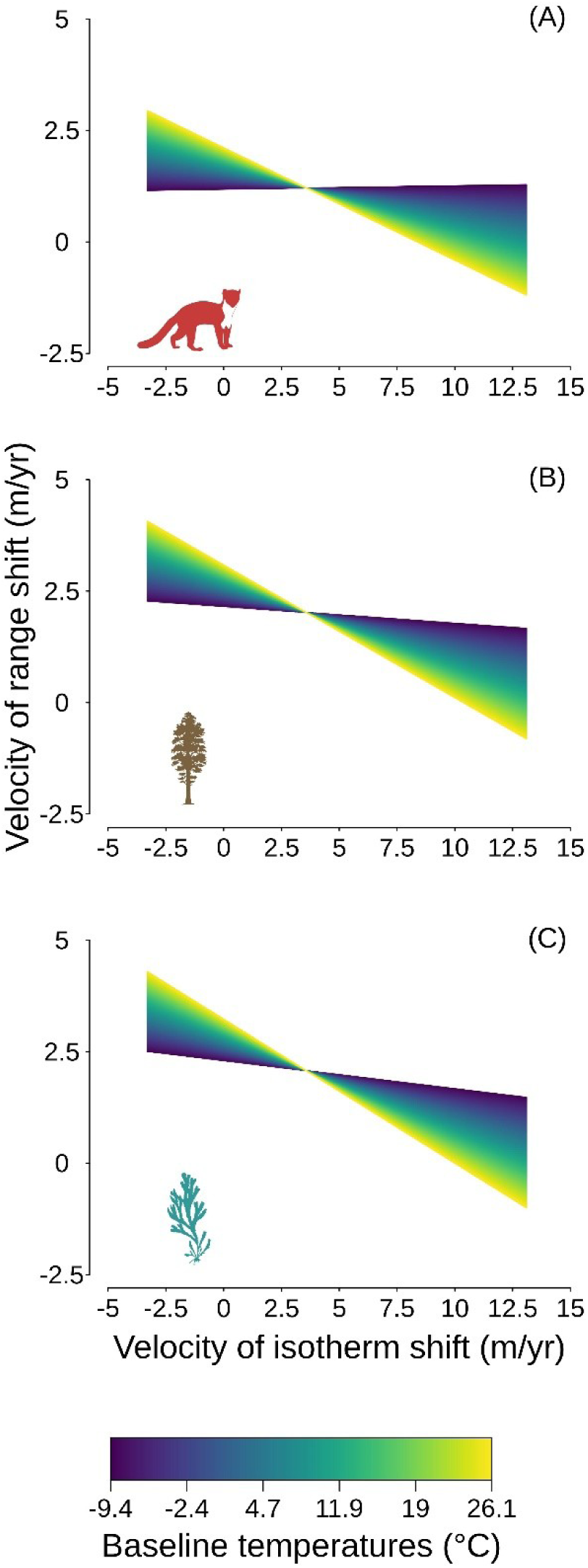
Interaction between mean annual temperature prior to the baseline survey (baseline temperatures) and the velocity of isotherm shifts in elevation for terrestrial (**A**) endotherms, (**B**) phanerogams and (**C**) cryptogams (see Fig. 3A for the case of ectotherms).

**Fig. S15.**
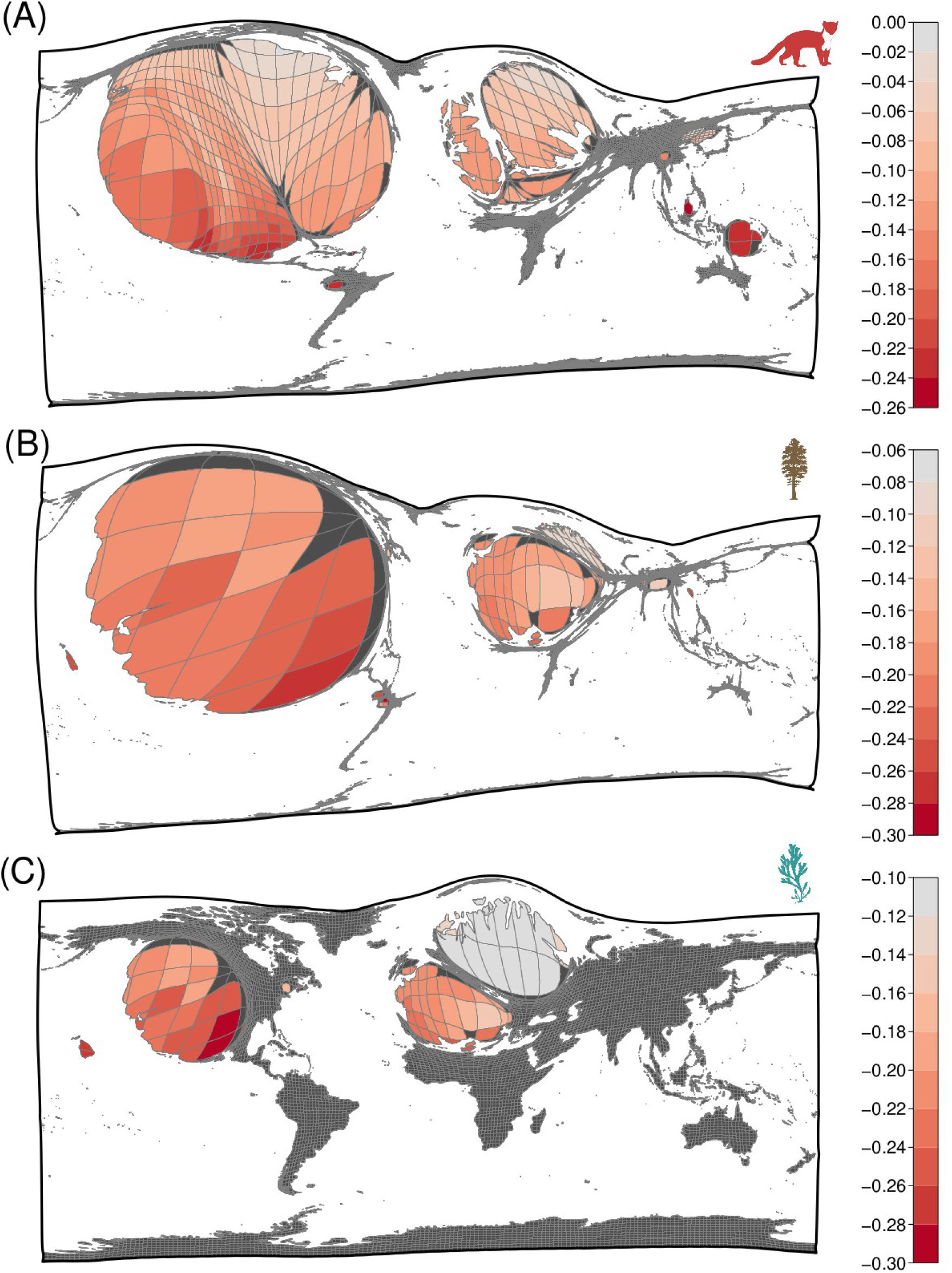
Cartograms of the predicted slope coefficient between the velocity of species’ range shifts and the velocity of isotherm shifts per 2° × 2° grid cell along the elevational gradient for (**A**) endotherms, (**B**) phanerogams and (**C**) cryptogams (see Fig. 4A for ectotherms). The number of range shift estimates (i.e. sample size) in each grid cell was used to distort the map: the bigger the grid cell, the larger the sample size.

## Supplementary Tables

**Table S1**. List of references from which the data were extracted.

**Table S2**. Full factorial design of spatial gradient (latitude *vs*. elevation) × positional parameter (centroid *vs*. margins) × biological systems (marine *vs*. terrestrial) × hemisphere (north *vs*. south) (N = 12 combinations). A total of 10 linear mixed-effects models and one linear model were calibrated to assess the mean rate of range shift per taxonomic class and at a given position within the range (centroid, leading edge and trailing edge). Model formulas are provided as used with the function “lmer” from the package “lme4” in the R programming language (28), except for the model of elevational range shifts at the margins of the distribution in terrestrial systems of the southern hemisphere for which there is no random effect structure. For this particular factorial model, the “lm” function was used instead. In total, 20 different taxonomic classes with more than 30 observations were considered in the analysis. Sample size per taxonomic class is given in parentheses. Summary statistics are provided for each model: the mean and standard deviation of the marginal and conditional R2 values across the 5,000 bootstrap iterations as well as the total contribution (%) of methodological variables which is the difference between the mean conditional and mean marginal R2 values.

**Table S3**. Summary statistics (mean, median, standard deviation and 95% confidence interval) of the estimated rate of range shift per taxonomic class for the different factorial models (see Table S2). A bootstrap approach based on 5,000 iterations was used to calculate the summary statistics for each of the estimated rate of range shift. Stars show significant effects (*: *P* < 0.05; **: *P* < 0.01; ***: *P* < 0.001) determined from the bootstrap distributions for the two alternative hypotheses of a mean rate of range shift being either negative or positive relative to a zero shift (i.e. the null hypothesis). The total sample size per taxonomic class, in terms of number of taxa, are also provided.

**Table S4**. Summary statistics (mean, median, standard deviation and 95% confidence interval) of the marginal and conditional R2 values obtained for each of the best candidate model relating the velocity of species range shifts against the velocity of isotherm shifts. A bootstrap approach based on 5,000 iterations was used to calculate the summary statistics.

**Table S5**. Outputs from the best candidate model relating the velocity of species range shifts against the velocity of isotherm shifts, either along the elevational (*EleVeloT*) or latitudinal (*LatVeloT*) gradient, while accounting for the effect of other covariates and their potential interaction with the velocity of isotherm shifts: baseline temperature conditions (*BaselineT*); the standardised human footprint index (*HFI*); and *LifeForm* (ectotherms, endotherms, cryptogams, phanerogams). A bootstrap approach based on 5,000 iterations was used to calculate the summary statistics (mean, median, standard deviation and 95% confidence interval) for each of the coefficient estimates. Stars show significant effects (*: *P* < 0.05; **: *P* < 0.01; ***: *P* < 0.001) determined from the bootstrap distributions for the two alternative hypotheses of a mean coefficient estimate being greater or lower than zero (i.e. the null hypothesis).

